# Entorhinal cortex directs learning-related changes in CA1 representations

**DOI:** 10.1101/2021.12.10.472158

**Authors:** Christine Grienberger, Jeffrey C. Magee

## Abstract

Learning-related changes in brain activity are thought to underlie adaptive behaviors^1,2^. For instance, the learning of a reward site by rodents requires the development of an over-representation of that location in the hippocampus^3-6^. However, how this learning-related change occurs remains unknown. Here we recorded hippocampal CA1 population activity as mice learned a reward location on a linear treadmill. Physiological and pharmacological evidence suggests that the adaptive over-representation required behavioral timescale synaptic plasticity (BTSP)^7^. BTSP is known to be driven by dendritic voltage signals that we hypothesized were initiated by input from entorhinal cortex layer 3 (EC3). Accordingly, the CA1 over-representation was largely removed by optogenetic inhibition of EC3 activity. Recordings from EC3 neurons revealed an activity pattern that could provide an instructive signal directing BTSP to generate the over-representation. Consistent with this function, exposure to a second environment possessing a prominent reward-predictive cue resulted in both EC3 activity and CA1 place field density that were more elevated at the cue than the reward. These data indicate that learning-related changes in the hippocampus are produced by synaptic plasticity directed by an instructive signal from the EC3 that appears to be specifically adapted to the behaviorally relevant features of the environment.

---

The behavioral experience of animals has been found to shape population activity in the hippocampus, and this experience-dependent neuronal activity is required to learn rewarded locations^3-6^. Such learning-related neuronal changes are commonly thought to be mediated by synaptic plasticity, generally of the Hebbian type^7-9^. To directly examine the physiological processes by which experience alters hippocampal population activity, we used two-photon Ca^2+^ imaging to record the activity of GCaMP6f-expressing CA1 pyramidal neurons^10^ in head-fixed mice engaged in a spatial learning task (Fig. 1a). The task consisted of two phases. Mice were first habituated to the linear track treadmill using a blank belt of 180 cm length, with the reward (10% sucrose/water) location varying from lap to lap (Fig. 1 b-k). On day 0 (final day of this habituation phase), the animals’ lick rates and running speeds were uniform throughout the environment (Fig. 1b-f), and CA1 place cells evenly tiled the space (Fig. 1g-h). In the second phase, the reward was delivered at a single fixed location, and the track contained several sensory cues uniformly distributed in space (day 1 = first exposure to the fixed reward location) (Fig. 1b-l). During this session, the animals gradually restricted their licking to the part of the environment around the reward (Fig. 1b-d) and concurrently slowed their running speed when approaching the reward delivery site (Fig. 1e-f). In parallel to these behavioral changes, we observed an increase in the total number of CA1 place cells, with the density of place cells near the reward site elevated over two-fold (Fig. 1g-i)^3,4,6^. Spatial information content (Fig. 1j) and lap-to-lap reliability (Fig. 1k) of individual place fields were also enhanced. Finally, the place cell population vector correlation was significantly lower when compared between days versus within days (Fig. 1k). Together, these results indicate that the learning of the reward location on day 1 is associated with an alteration in the CA1 representation that includes a strongly elevated place cell density near the reward, the presence of which is significantly correlated with low running speeds measured around the rewarded location (Fig. 1l). This so-called reward over-representation is similar to the CA1 activity adaptations previously found to be required for the successful learning of the reward location^5^.

**Figure 1:**
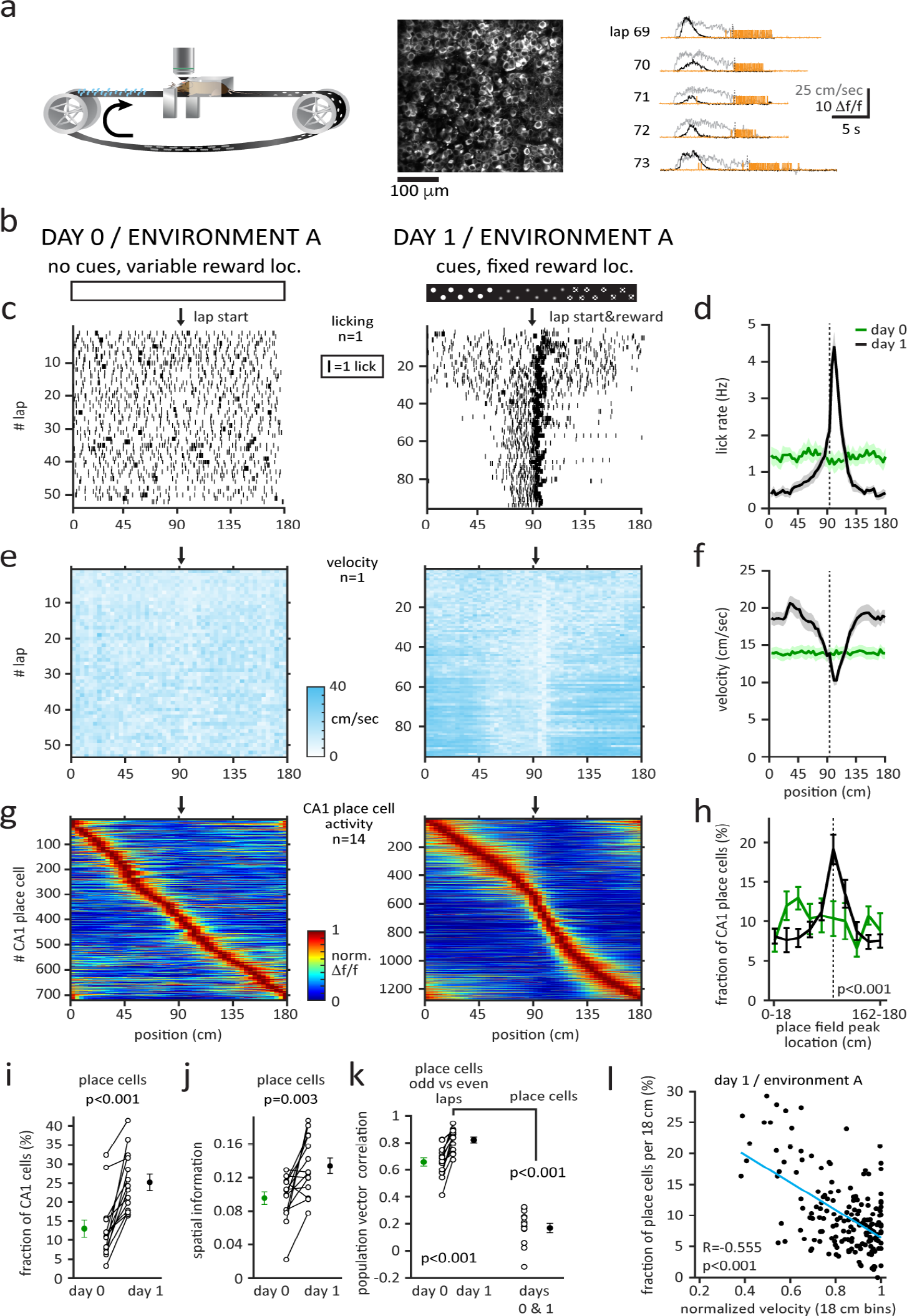
Experience-dependent shaping of area CA1 representations. **a**, Left. Diagram of the experimental setup. Mice are trained to run on a linear treadmill for a sucrose reward. Middle: Two-photon, time-averaged image showing GCaMP6f expression in CA1 pyramidal neurons. Right: Ca^2+^ Δf/f traces (black) for five consecutive laps from a CA1 place cell. Simultaneously recorded velocity and licking signals are shown in grey and orange. **b**, Two task phases (environment A). Left: The training and one recording session (day 0) are performed on a blank belt with from lap-to-lap varying reward locations. Right: Next (day 1), the mouse is exposed to a sensory cue-enriched belt and must learn the fixed reward location. **c**, Licking behavior of an individual animal. The ticks represent licks; the arrows mark the lap start (left) or lap start/reward location (right). **d**, Mean lick rates for days 0 (green) and 1 (black) for n=18 animals. **e**, Running behavior of an individual animal. **f**, Mean running for days 0 (green) and 1 (black). **g**, Normalized mean Δf/f across space for all CA1 place cells (PCs) (day 0: n=719, day 1: n=1278). PCs are ordered according to their peak location on the track. Only data from animals with the same field of view imaged in both sessions (n=14) are included. **h**, Fraction of CA1 PCs as a function of place field peak location (day 0: green, day 1: black). The track is divided into ten spatial bins of 18 cm each (chi-square test, p<0.001). **i**, Fraction of CA1 cells that are spatially modulated (paired two-sided *t*-test, p<0.001). **j**, Mean PC spatial information per animal (paired two-tailed *t*-test, p=0.003). **k**, Population vector (PV) correlations. Left. Reliability of CA1 PC activity. Place cell PVs for odd and even laps were correlated (paired two-tailed *t*-test, p<0.001). Right: PV correlations for CA1 cells with place fields on days 0 and 1 (two-tailed *t*-test, p<0.001). **l**, CA1 PC density as a function of the normalized velocity. Each dot represents 18 cm. Data pooled across all animals (n=18) and fit by linear equation (blue line). In panels i-k, the open circles show individual animals, the filled circles the mean. Black dashed lines/arrows mark the reward location. Data are shown as mean +/- SEM.

Next, we asked whether a recently discovered synaptic plasticity type, behavioral timescale synaptic plasticity (BTSP)^11^, could underlie the experience-dependent formation of the CA1 representation on day 1. BTSP is exclusively driven by long-duration dendritic voltage signals, Ca^2+^ plateau potentials (“plateaus”), that can induce plasticity in a single trial (Fig. 2a, left and middle)^12-16^. Moreover, BTSP follows an asymmetric learning rule that operates on the behaviorally relevant time scale of seconds. Therefore, it produces predictive neuronal activity, that is, the firing field generated by BTSP precedes the plateau in space by an amount that depends on the running speed (Fig. 2a, left and middle). Another effect of the BTSP timescale is that there is a direct relationship between the animal’s running speed in the lap where the place field first appeared (‘induction lap’) and the width of the resulting place field in space (Fig. 2a, right). Consistent with the involvement of this new type of synaptic plasticity, we observed that previously silent neurons acquired place fields abruptly in a single lap (Fig. 2b), with a substantial fraction of new place fields added during the learning session (Fig. 2c-d, Extended Data Fig. 1). These suddenly appearing place fields exhibited additional hallmark features of BTSP^11^. First, place fields tended to shift backward in space compared to the activity in their induction lap (Fig. 2e). Second, we observed a linear relationship between the place field’s width and the animal’s velocity in the induction trial (Fig. 2f). Finally, the development of the experience-dependent representation during the session, including the reward overrepresentation, was significantly inhibited by local application of a pharmacological antagonist of synaptic plasticity, D-APV (Fig. 2g), or an inhibitor of plateau firing, the CaV2.3 channel blocker SNX-482 (Fig. 2h). Presumably, the local nature of the antagonist application limited the behavioral impact of the manipulation as all behavioral measures were unaltered (Extended Data Figs. 2-3). The above results suggest that the experience-dependent shaping of the CA1 representation requires BTSP.

**Figure 2:**
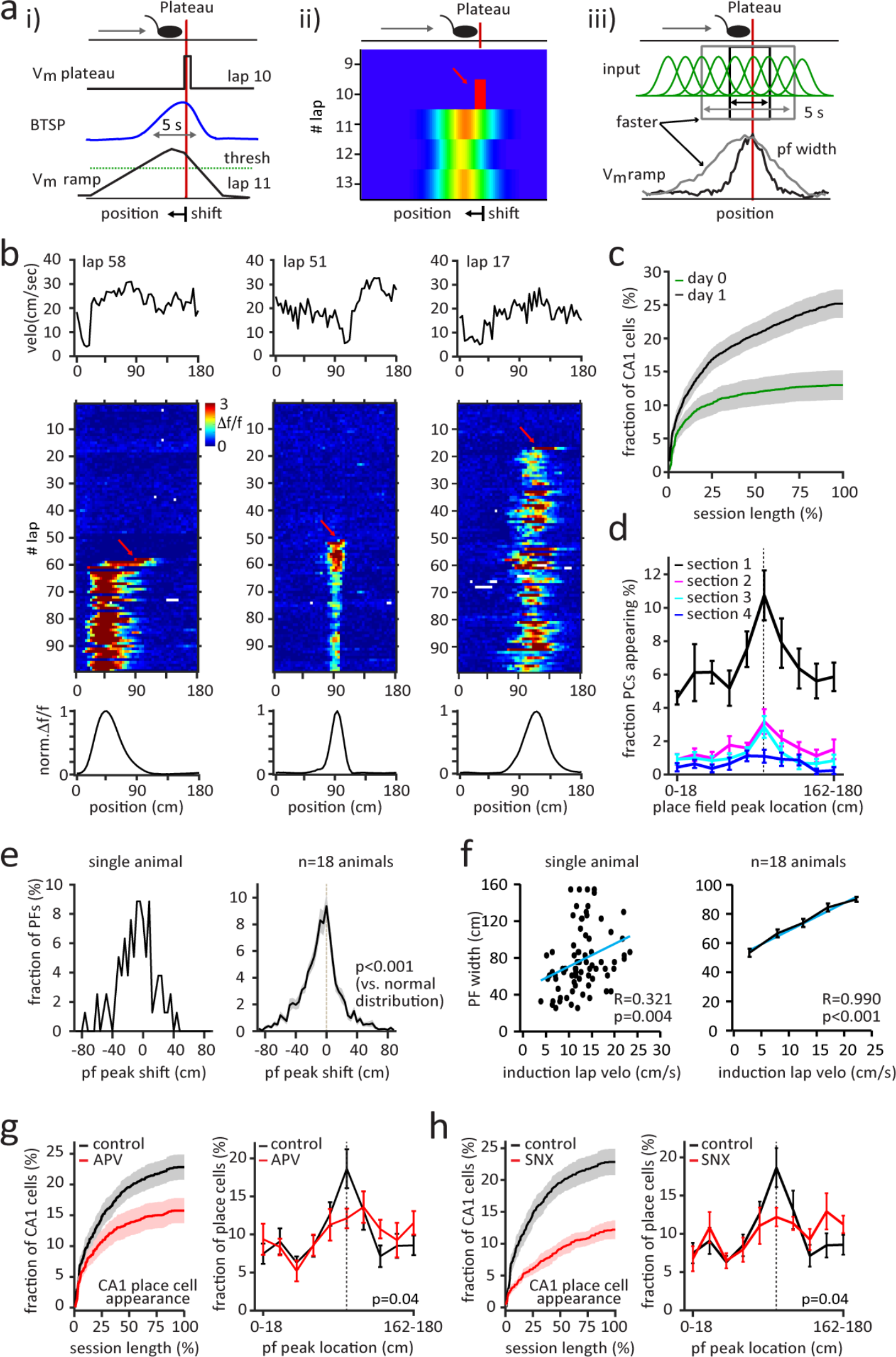
Behavioral timescale synaptic plasticity underlies experience-driven shaping of CA1 representations. **a**, Schematics showing features of behavioral timescale synaptic plasticity (BTSP). **b**, Three abruptly appearing place cells (PCs). Top: velocity profile of the first lap with place field activity (= induction lap). Middle: Δf/f across laps. The red arrows mark the induction lap. Bottom. Normalized mean Δf/f across space. **c**, Time course of CA1 PC appearance for days 0 (green) and 1 (black). **d**, Fraction of CA1 PCs as a function of place field peak location and session length (session divided into four sections of 14-35 laps). **e**, Histograms showing place field (pf) peak shift (peak location (cm) of the generated place field minus peak activity location (cm) in the induction lap). Left: individual animal. Right: n=18 animals (n=1727 CA1 place cells, one-sample Kolmogorov-Smirnov test, p<0.001). **f**, Place field width as a function of the animal’s mean induction lap velocity. Left: individual animal. Each dot represents one place cell. Right: n=18 animals. Data is binned into 5 cm/sec velocity bins and fit by linear equation (blue line). **g**, Effect of NMDA receptor antagonist, D-APV (50/75 µM), on the development of the CA1 representations. Black: Control (n=10 animals). Red: APV (n=8 animals). Left, Time course of CA1 PC appearance. Right: Fraction of CA1 PCs as a function of pf peak location (chi-square test, p=0.04). **h**, Effect of Ca^2+^ channel antagonist, SNX-482 (10 µM). Black: Control (n=10 animals). Red: SNX (n=7 animals). Panels same as **g** (chi-square test, p=0.04). Black dashed lines depict the reward location. Data are shown as mean +/- SEM.

EC3 axons innervate the apical dendritic tuft^17-19^, which is the site of plateau initiation in CA1, and previous *in vitro* and *in vivo* work identified a crucial role of the EC3 input in driving plateaus^12,15^. Therefore, we next asked if perturbing EC3 input could impact the formation of the CA1 over-representation. To examine this, we used a retrograde virus infection strategy^20^ to express the hyperpolarizing proton pump Archaerhodopsin-T (ArchT)^21^ only in the subset of EC3 neurons that projected to our recording area in CA1 (Fig. 3a). As a control, we expressed tdtomato instead of ArchT in a separate group of mice that, otherwise, received the same treatment. We found that inhibiting this subset of EC3 axons by delivering 594 nm laser light (40 Hz, sinusoidal stimulation)^22^ to the EC in a zone of 36 cm (+/- 18 cm) around the reward prevented the development of the CA1 reward overrepresentation compared to the control group (Fig. 3b-c, Extended Data Fig. 4). Notably, there was no significant change in the amplitude of place fields near the reward zone (n=6 mice (tdtomato) vs. n=8 mice (ArchT), 78% vs. 84 % Δf/f, two-tailed unpaired *t*-test, p=0.401), mean Ca^2+^ event amplitude (Extended Data Fig. 4c, n=6 mice (tdtomato) vs. n=8 mice (ArchT), two-tailed unpaired *t*-test, p=0.06), the time course of place field formation (Extended Data Fig. 4d) or in the licking and running behaviors (Extended Data Fig. 4e) between the control mice and the ArchT group.

**Figure 3:**
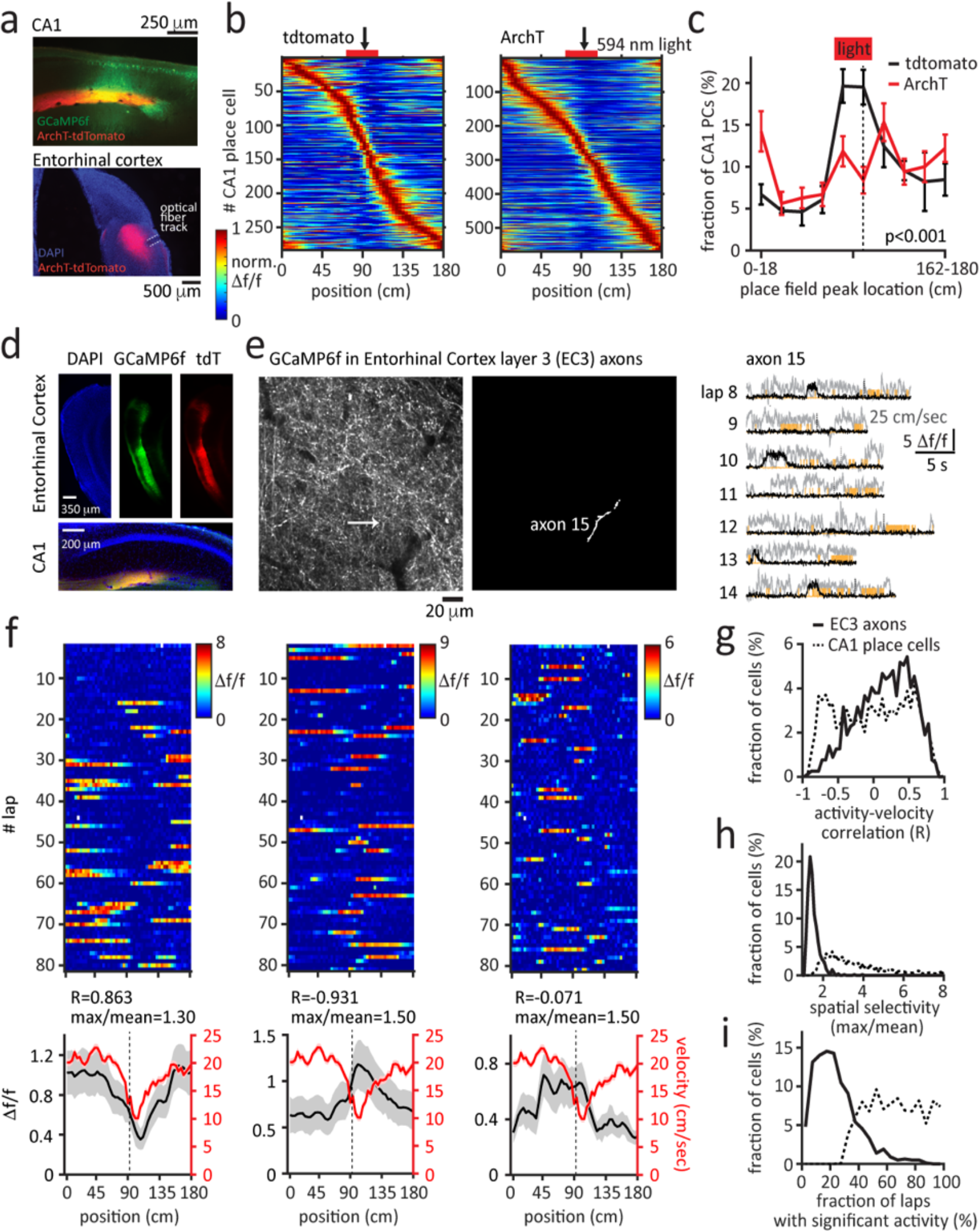
Entorhinal cortex layer 3 activity is required for the experience-dependent shaping of CA1 representations. **a-c**, Ipsilateral optogenetic perturbation of entorhinal cortex layer 3 (EC3) neuronal activity prohibits the development of the reward overrepresentation. Black: tdtomato control (n=6 animals), Red: ArchT (n=8 animals). **a**, Viral expression of GCaMP6f in CA1 (top) and ArchT-tdtomato in EC3 (bottom). EC3 axons can be found in the stratum lacunosum-moleculare (SLM) of CA1 (top). **b**, Normalized mean Δf/f across space for all CA1 place cells (PCs). **c**, Fraction of CA1 PCs as a function of place field peak location (chi-square test, p<0.001). **d-i**, Recording of EC3 axonal activity in CA1 SLM. **d**, Viral expression of GCaMP6f and tdtomato in EC3 neurons (top) and their axons in hippocampal area CA1 (bottom). **e**, Left. Two-photon, time-averaged image showing expression of GCaMP6f in EC3 axons. Middle: white area depicting axon 15. Right: Ca^2+^ Δf/f traces (black) for 7 consecutive laps recorded from axon 15. Simultaneously recorded velocity and licking signals are shown in grey and orange. **f**, Three individual EC3 axons. Top: Δf/f across laps. Bottom. Mean Δf/f (black) and mean velocity (red) across space. Black and red y-axes apply, respectively. EC3 axons are classified based on their mean Δf/f-mean velocity-correlation (Pearson correlation, R) and spatial selectivity index (max Δf/f divided by mean Δf/f). **g-i**, Distributions of the activity-velocity correlations (**g**), the spatial selectivity indices (**h**), and the fractions of laps with significant activity (**i**) from 792 axons from 7 animals (solid line) and 1727 CA1 PCs from 18 animals (dashed line). Black arrows and dashed lines depict the reward location. Data are shown as mean +/- SEM.

As EC3 appears to be necessary for the experience-dependent shaping of the CA1 representation, we next wanted to examine the activity of EC3 neurons projecting to CA1. The axons of these cells are located in the stratum lacunosum-moleculare of CA1 (Fig. 3d) and are accessible for two-photon imaging through our standard hippocampal window. Thus, we performed axonal two-photon Ca^2+^ imaging in mice expressing GCaMP6f in EC3 neurons (Fig. 3e, Extended Data Fig. 5). Evidence for selective activity in the individual axons was observed as spatial (average Δf/f max/mean) and velocity tuning (Pearson correlation coefficient between average Δf/f and average velocity) (Fig. 3f-i) ^23-26^. However, a large amount of stochastic variation was obvious in both the spatial and velocity tuning, when the average axon Δf/f was compared for interleaving trials (median Pearson’s correlation coefficient = 0.247, odd vs. even trials, Fig. 4a-b). Only a fraction of axons (19%) showed relatively well-correlated firing (peak locations on even trials were within 10 cm of odd trials), and the peak firing locations of these well-correlated axons uniformly tiled the entire track (Extended Data Fig. 6a-c). Similar results were seen for the 5% most selective EC3 axons (Extended Data Fig. 6d-f)^24^. These data indicate that EC3 activity during this behavior is largely stochastic with a small degree of structure mainly provided by a subset of axons, whose peak activity uniformly tiled the environment.

**Figure 4:**
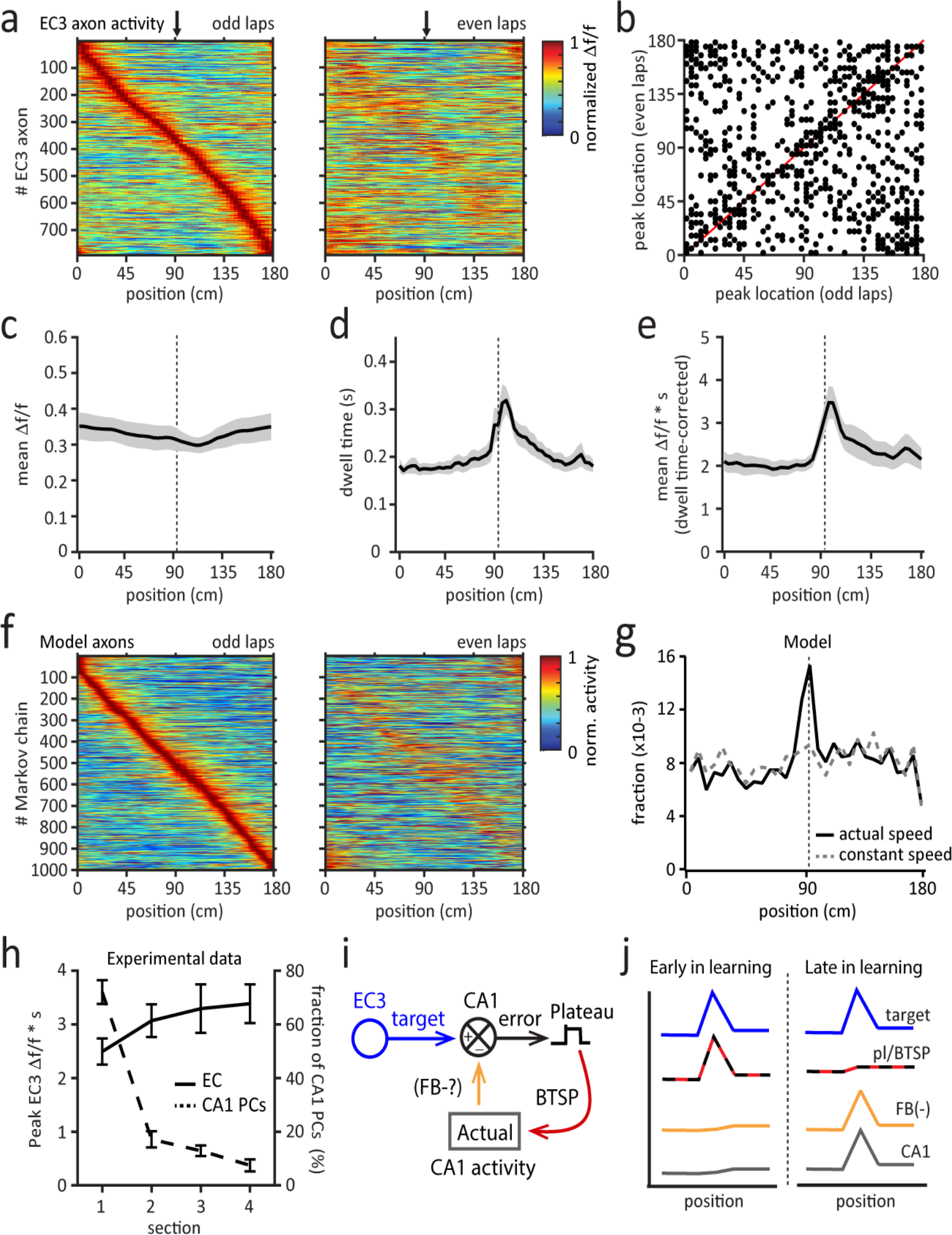
Entorhinal cortex layer 3 generates a target. **a**, Normalized mean Δf/f across space for all entorhinal cortex layer 3 (EC3) axons recorded (n=792 axons in 7 animals). Color plots for odd and even laps are shown separately. EC3 axons are ordered according to their peak location during the odd laps. **b**, Scatter plot showing axonal activity peak locations for averages made from odd and even laps. The unity line is depicted in red. **c**, Mean Δf/f activity across all EC3 axons and all laps. **d**, Mean dwell time (s) across space. **e**, Occupancy-corrected EC3 axon activity, represented by the integrated Ca^2+^ signals (mean Δf/f *s). **f**, Same as panel **a** for 1000 modeled EC3 axons. **g**, Model predicts the fraction of postsynaptic neurons crossing threshold for plateau initiation across space. Black: experimentally observed running profile. Grey: Constant running speed. **h**, Mean occupancy - corrected EC3 axon activity around the reward (solid black line) and the fraction of CA1 place cells appearing near the reward (dashed black line) as a function of session length. The session is divided into four sections of 14-35 laps. **i**, EC3 provides each individual CA1 neuron with a desired or target activity pattern that is compared in the distal apical dendrites with a representation of the actual pattern of population activity that could be provided by feedback via OLM-Interneurons (FB-?). The plateau functions as a local error signal in each CA1 cell that drives BTSP and shapes the firing of each CA1 cell accordingly. **j**, Spatial profiles of hypothesized signals. Black arrows and dashed lines depict the reward location. Data are shown as mean +/- SEM.

As expected from the above analysis, the mean Δf/f for all axons was basically constant across the linear track’s spatial locations (Fig. 4c). Hence, after factoring in the animals’ spatial running profile (Fig. 4d), the occupancy-corrected EC3 activity (i.e., Δf/f multiplied by spatial bin dwell time) showed a reliable, approximately two-fold, increase near the reward (Fig. 4e). These results suggest that, as a population, EC3 axons exhibit a constant firing rate (AP/s) across the track that increases the number of APs occurring around the reward area simply because the animals dwell there for a longer amount of time.

To understand how this EC3 activity pattern could impact postsynaptic CA1 pyramidal neurons, we turned to computational modeling (Extended Data Fig. 6f-i). Because EC3 single axon activity was suggestive of a stochastic process (e.g., exponentially distributed activity times and low cell-cell correlations, Fig. 4a-b, Extended Data Figs. 6a-c, i), we modeled the activity of individual EC3 axons as simple two-state Markov chains (Extended Data Fig. 6f-i). This approach simulates EC3 neuron activity transitions between an inactive and active state, similar to the persistent firing previously observed in this region^26-29^. While a completely homogeneous version of this model, without any modulation of the transition probabilities, recapitulated many aspects of the EC3 data (i.e., selectivity index, velocity correlation, uniformly distributed peak activities), it did not show the small subset of well-correlated activity that tiled the environment (Extended Data Fig. 6j-o). To produce this uniformly distributed set of cell-cell correlations, it was necessary to adjust the activation transition probability uniformly across the track (P^01^ increased from 0.04 to 0.24 for 10 time steps at a point in the lap that incremented smoothly across the population; Fig. 4f, Extended Data Fig. 7a-f).

We next used the model to calculate the probability that a given CA1 postsynaptic neuron receives a suprathreshold, plateau-evoking, amount of EC3 input (Extended Data Fig. 7g-i). The simulation predicts that the constant level of EC3 input across the track produces a steady probability of plateau potential initiation (dashed line in Fig 4g, assuming a constant running speed). Therefore, when using the actual running behavior of the mice (solid line in Fig. 4g), the fraction of neurons initiating plateaus at the reward site increases approximately two-fold, again, because the animals spend approximately twice the time at this location. Taken together, the EC3 recordings and the simple model indicate that the EC3 has produced a pattern of activity that reflects the uniformity of the environment and uses the running behavior of the animal to increase plateau initiation probability and therefore BTSP induction near the reward site.

The above data suggest that EC3 input may direct neuronal plasticity in CA1 by providing a type of instructive signal. Therefore, we next attempted to determine the form of this EC3 instructive signal. If functioning as an error signal, we would expect EC3 activity around the reward site to decrease as CA1 population activity approached the desired pattern. On the other hand, if the EC3 provides a signal representing the desired CA1 activity pattern (a target signal) to each CA1 pyramidal neuron, it should remain more constant throughout the session, even as CA1 plasticity decreases^30^. To examine this, we plotted EC3 population activity as a function of the session duration alongside the plasticity of CA1 place cells. We found that, while the formation of new CA1 place cells decreased markedly during the session, the EC3 activity profile remained elevated throughout the entire session (Fig. 2b-c, Fig.4h). Altogether, these results suggest that the EC3 provides an invariant instructive signal that is more reminiscent of a target signal than an error signal. In our current scheme, the excitatory target from EC3 combines in the apical dendrites with an inhibitory signal representing the actual CA1 population activity, and the resulting plateau potentials function as a local error signal that is unique in each CA1 neuron^31,32^ (Fig. 4i). The source of the actual CA1 activity signal remains undetermined and may involve inhibitory, neuromodulatory, disinhibitory or other elements^5,19,33-36^. The hypothesized evolution of these signals during a learning session is schematized in Fig. 4j.

Finally, we tested how the EC3 target signal responds to a less uniform environment that contains only a single prominent, novel, and reward-predictive feature. To do so, we designed a different environment (environment B) that included a visual stimulus (500 ms-long blue light flashes (10 Hz) to both eyes) activated 50 cm before the fixed reward delivery site and no other experimenter-placed belt cues (Fig. 5a, red). This second environment elicited substantial changes in the running of the mice (Fig. 5c) and more subtle alterations in their licking (Fig. 5b). In addition, the EC3 axon population activity was substantially altered. The most prominent change was an approximately 3-fold increase in the fraction of axons whose firing peaked around the visual stimulus (Figure 5d, Extended Data Fig. 8a-c). Nevertheless, EC3 axon activity retained a high degree of instability with only a fraction of axons (25%) showing relatively consistent firing between interleaved trials (Fig. 5d-e, Extended Data Fig. 8 d-e). Interestingly, the density of these well-correlated axons was enriched at the location of the light stimulus (Fig 5e, Extended Data Fig. 8d), indicating that the activity of the EC3 neurons reflected the distribution of relevant environmental cues. To reflect this new EC3 activity data more accurately, we adjusted the previous Markov chain simulation by increasing the fraction of chains that had an elevated activation probability around the visual cue position such that the final spatial profile of peak chain activity showed an approximately 3-fold increase around this position (see methods, Figure 5f; Extended Data Fig. 8f-g). Moreover, this modification recapitulated the increased density of the well-correlated axons around the visual stimulation location (Fig. 5f-g, Extended Data Fig. 8h-i). When we ran the simulation using this set of parameters to infer plateau probability across space, our results predicted the presence of an over-representation of the visual stimulus location that was even larger than that produced by the running behavior alone (Fig. 5h; Extended Data Fig. 8j).

**Figure 5:**
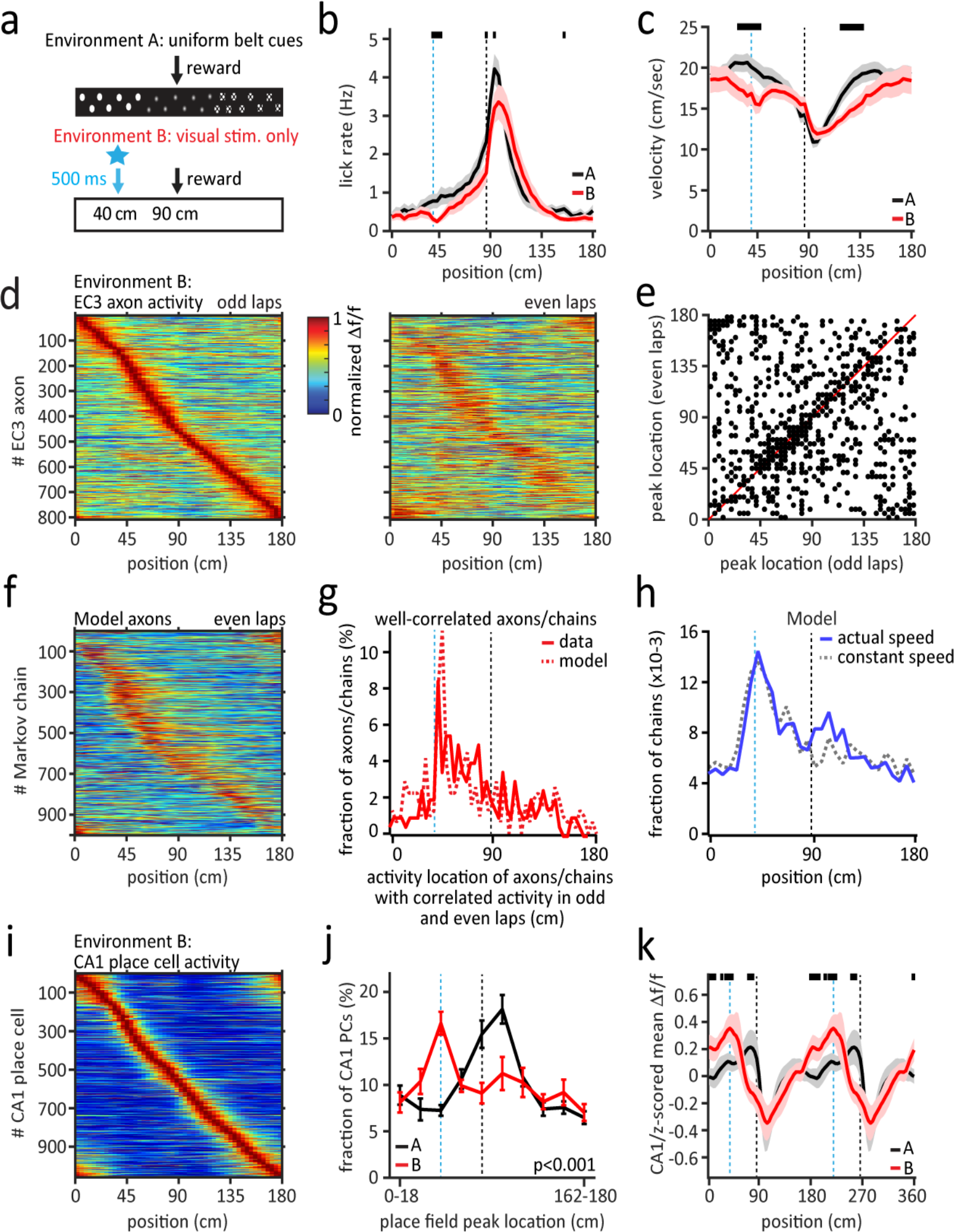
The entorhinal cortex layer 3 target signal adapts to the environment. **a**, In contrast to environment (env) A (black), env B (red) involves a blank belt and a visual stimulus (blue LED flashes, 10 Hz, 500 ms) 50 cm before the single, fixed, reward (black arrow). **b**, Mean lick rate in env A (black, n=25 mice) and B (red, n=17 mice). **c**, Mean running profile in env A (black) and B (red). **d**, Normalized mean Δf/f across space for EC3 axons (n=808, n=8 animals) in env B. Color plots for odd and even laps are shown separately. EC3 axons are ordered according to their peak location during the odd laps. **e**, Scatter plot showing axonal activity peak locations for averages made from odd and even laps. The unity line is depicted in red. **f**, Normalized mean Δf/f across space for modeled Markov chains. **g**, Histogram of the peak activity locations of well-correlated axons (red, solid, 25% of all EC3 axons) and chains (red, dashed, 31% of all chains). Peak locations on even trials were within 10 cm of odd trials. **h**, Model predicts the fraction of postsynaptic neurons crossing plateau initiation threshold across space (blue, solid: experimentally observed running profile; grey, dashed: constant running speed). **i**, Normalized mean Δf/f across space for all CA1 place cells (PCs) (n=1058, n=9 animals) in env B. **j**, Fraction of CA1 PCs as a function of place field peak location (A: n=18, black; B: n=9, red; chi-square test, p<0.001). **k**, z-scored mean Δf/f of all CA1 PCs. Two identical mean activity profiles are concatenated. Black bars in panels b, c, l indicate locations with p<0.05 (unpaired two-tailed *t*-test, performed on each spatial bin). Blue dashed lines depict the light onset, black dashed lines the reward location. Data are shown as mean +/- SEM.

We tested this prediction by performing Ca^2+^ imaging in CA1 pyramidal cells in mice exposed to environment B. We first found that the basic characteristics of place cell activity, such as the fraction of spatially modulated cells and the place cell spatial information content per animal, remained unchanged (Extended Data Fig. 9). However, consistent with our modeling data, we observed the largest fraction of place cells near the location of the light (Fig. 5i-j) and a ramping mean Δf/f activity profile that peaked near the visual stimulus in environment B rather than near the reward site as in environment A (Fig. 5k). Accordingly, the running speed profile is a less powerful predictor of the reward over-representation location in environment B than in environment A (Extended Data Fig. 10). These results corroborate our hypothesis that EC3 is necessary and sufficient for shaping the CA1 representation. In fact, EC3 provides a target signal that instructs CA1 in how to represent the environment during a spatial learning task. Furthermore, because the EC3 activity changed from a spatially uniform pattern in environment A to one with enhanced activity at the newly introduced reward-predictive feature in environment B, it appears that the entorhinal cortex is sensitive to behaviorally relevant aspects of an environment, and perhaps self-motion cues.

This work addresses the long-standing question of what neural mechanisms underlie learning within the mammalian brain (Extended Data Fig. 11). Our results show that a recently discovered synaptic plasticity (BTSP) is responsible for the adaptive changes that occur in hippocampal area CA1 population activity as an animal learns a simple task. We also present several lines of evidence that the EC3 is the source of a target-like instructive signal that directs this plasticity to achieve a desired CA1 population activity. Finally, stochastic EC3 activity was observed to appropriately adapt to the distribution of salient cues in the environment, which ranged in uniformity. While the stochasticity observed in EC3 axon activity is suggestive of certain computations (e.g., sampling methods for probabilistic inference)^37,38^, additional experiments are required to confirm this mode of activity and to determine exactly how entorhinal cortex neurons are adjusted to produce an environmentally specific instructive signal.

Target signals can theoretically be quite powerful in directing learning in complex neuronal networks because they provide a means to account for the multitude of downstream parameters that lie between regional brain activity and desired behavior^39,40^. However, reports of target signals driving synaptic plasticity, Hebbian or otherwise, are rare^30^. Indeed, even brain regions thought to employ supervised motor learning have been found to use error signals, not targets^41,42^. The observation that a synaptic plasticity directed by adapting target signals shapes the activity of the mammalian hippocampus, an area well-known for its importance in spatial learning and episodic memory, raises the possibility that many brain regions may learn in a manner substantially different than currently thought.

## Acknowledgements

We thank Randy Chitwood for technical assistance. We thank Randy Chitwood, Sandro Romani, and Nelson Spruston for useful discussions. This work was supported by the Howard Hughes Medical Institute.

## Author Contributions

CG and JCM designed research. CG performed *in vivo* recordings. CG and JCM analyzed the experimental data. JCM designed and implemented the computational model. CG and JCM wrote the manuscript.

## Online methods

All experiments were performed according to methods approved by the Janelia (Protocol 12-84 & 15-126) and the Baylor College of Medicine’s (Protocol AN-7734) IACUC committees.

### Surgery

All experiments were performed in adult (older than P66 at the time of surgery) GP5.17^10^ (n=52 mice, Janelia and Jackson Laboratories) or pOxr1-Cre^19^ (n=34 mice, Jackson Laboratories) mice of either sex by an experimenter who was not blind to the experimental conditions. Animals were housed under an inverse 12-hour dark/12-hour light cycle. All surgical procedures were performed under deep isoflurane anesthesia. After locally applying topical anesthetics, the scalp was removed, and the skull was cleaned. Then the skull was leveled, and the locations for the craniotomies were marked using the following stereotactic coordinates: 1) center of the 3mm-diameter hippocampal window: 2.0 mm posterior from Bregma and 2.0 mm lateral from the midline; 2) CA1 virus injections: 2.0 mm posterior from Bregma and 1.9 mm lateral from the midline and 2.3 mm posterior from Bregma and 2.2 mm lateral from midline; 3) EC virus injections: 4.7 mm posterior from Bregma and 3.5 mm from the midline and 4.7 mm posterior from Bregma and 3.8 mm from the midline; 4) entorhinal cortex (EC) optical fiber implantation: 4.7 mm posterior from Bregma and 4.4 mm from the midline; and 5) EC local field potential (LFP) recordings: 4.7 mm posterior from Bregma and 3.5 mm from the midline. Then, for all experiments except the LFP recordings, a 3 mm-diameter craniotomy was made above the hippocampus. Cortical tissue within the craniotomy was slowly aspirated under repeated irrigation with warmed sterile saline 0.9%. Once the external capsule was exposed, the cannula (3 mm diameter, 1.7 mm height) with a window (CS-3R, Warner Instruments) on the bottom was inserted and cemented to the skull. Finally, a custom-made titanium head bar was attached to the skull using dental acrylic (Ortho-Jet, Lang Dental).

For the experiments with GCaMP6f and tdtomato expression in entorhinal cortex layer 3 (EC3) or Archaerhodopsin-T (ArchT) or tdtomato expression in EC3 and GCaMP6f expression in CA1, the hippocampal window surgery was preceded in the pOxr1-Cre mice^19^ (n=34) by ipsilateral virus injections using the coordinates stated above. Notably, the pOxr1-Cre mouse line expresses Cre recombinase predominantly in the medial entorhinal cortex. For the virus injections, we first made a small (∼0.5 mm diameter) craniotomy. This was followed by injecting a small volume of one of the following mixtures (all viruses produced by the Janelia Viral Vector Core; viral titers range between 1 and 7.5E-12): 1) AAV1.Syn.GCaMP6f.WPRE.SV40 and AAVrg.Syn.Flex.ArchT-tdtomato.WPRE.SV40 into area CA1 (DV: 1350 and 1000 µm; 25 nl per depth); 2) AAV1.Syn.GCaMP6f.WPRE.SV40 and AAVrg.Syn.Flex.tdtomato.WPRE.SV40 into area CA1 (DV: 1350 and 1000 µm; 25 nl per depth); 3) AAV1.Syn.Flex.GCaMP6f.WPRE.SV40 and AAV1.Syn.Flex.tdtomato.T2A.tdtomato.WPRE.SV40 into the entorhinal cortex (DV: 2100, 1800, and 1500 µm; 50 nl per depth). The injection system comprised a pulled glass pipette (broken and beveled to 15–20 μm (outside diameter); Drummond, Wiretrol II Capillary Microdispenser), backfilled with mineral oil (Sigma). A fitted plunger was inserted into the pipette and advanced to displace the contents using a manipulator (Drummond, Nanoject II). Retraction of the plunger was used to load the pipette with the virus. The injection pipette was positioned with a Sutter MP-285 manipulator. For the optogenetics experiments, an optical fiber (core diameter of 200 µm) was chronically implanted at a 45° angle into the ipsilateral entorhinal cortex (at a depth of 50-100 µm) and attached to the skull using dental cement (Calibra Dual Cure, Pearson Dental).

### Behavioral training and task on the linear track treadmill

The linear track treadmill consisted of a belt made from velvet fabric (McMaster Carr). The belt (length of 180 cm) was self-propelled, and the reward was delivered through a custom-made lick port controlled by a solenoid valve (Parker). The animal’s speed was measured using an encoder attached to one of the wheel axles. A microprocessor-based (Arduino) behavioral control system interfaced with a Matlab GUI controlled the valve, the sensors, and the encoder. In addition, a separate microprocessor (Arduino) interfaced with a Matlab GUI was used to operate the laser shutter for the optogenetic perturbation experiments and control the visual stimulation based on the animal’s position on the belt. Behavioral data was monitored and recorded via a PCIe-6343, X series DAQ system (National Instruments), and the Wavesurfer software (Janelia).

5-7 days after the optical window implantation running wheels were added to the home cages, and mice were placed on water restriction (1.5 ml/day). After both training and recording sessions, mice were supplemented with additional water to guarantee a 1.5 ml/day water intake. After 5-6 days of familiarizing the animals with the experimenter, mice were trained to run head-fixed on the linear treadmill for 3-5 days. This training was conducted during the animals’ dark cycle, and mice were trained on a blank belt (no sensory cues) to run for a 10% sucrose/water reward delivered at from lap-to-lap varying locations.

To record neuronal activity and study the development of CA1 representations as mice learned to navigate in a new environment, we exposed the animals to two different environments (‘day 1’). Environment A consisted of a belt enriched with three different visual and tactile cues (glue sticks, Velcro tape patches, white dots), which covered the entire length of the belt^11,12,14^. The reward was delivered at a single, fixed reward location. For environment B, the belt was devoid of any local cues, and a bilateral visual stimulus (blue LED positioned in front of both eyes, flashing at 10 Hz for 500 ms) was delivered 50 cm before the fixed reward location. Individual recording sessions lasted between 45 and 60 min, with one recording session per day.

### In vivo two-photon Ca^2+^ imaging

All Ca^2+^ imaging recordings were performed in the dark using a custom-made two-photon microscope (Janelia MIMMS design). GCaMP6f and, if expressed, tdtomato were excited at 920 nm (typically 40 – 70 mW) by a Ti:Sapphire laser (Chameleon Ultra II, Coherent) and imaged through a Nikon 16x, 0.8 Numerical Aperture (NA) objective. Emission light passed through a 565 DCXR dichroic filter (Chroma) and either a 531/46 nm (GCaMP channel, Semrock) or a 612/69 nm (tdtomato channel, Semrock) bandpass filter. It was detected by two GaAsP photomultiplier tubes (11706P-40SEL, Hamamatsu). Images (512 × 512 pixels) were acquired at ∼30 Hz using the ScanImage software (Vidrio).

Specifics of CA1 pyramidal neuron Ca^2+^ imaging: imaging fields (size varied from 280 × 280 to 380 × 380 µm) were selected based on the presence of Ca^2+^ transients in the somata. One field of view was imaged per day. If possible, the same field of view was imaged on days 0 and 1 (n=14/18 animals).

Specifics of EC3 axonal Ca^2+^ imaging: imaging fields (size of 230 × 230 µm) were selected based on the presence of the fiber morphology in the tdtomato channel and the occasional Ca^2+^ transient in the field of view. No attempt was made to locate the same imaging field from day to day.

### Local pharmacology during two-photon imaging

For the local pharmacology experiments, the animal was briefly anesthetized ∼45 minutes before the recording session using isoflurane. Then the hippocampal window was carefully punctured (∼50-100 µm-wide hole) near the imaging field of view. This procedure lasted about 5-10 min. In the case of the APV experiments, the hole was then covered with a silicone elastomer (Kwik-Cast, wpi), and the animal was allowed to recover from the anesthesia for ∼45 minutes. Then, after positioning the animal under the two-photon microscope, we removed the Kwik-Cast plug and filled the cannula either with D-APV (50-75 µM) dissolved in sterile saline or with sterile saline alone. The animal was prevented from running for the initial 5-10 min to allow for the initial diffusion of the drug. APV continued to be present in the cannula throughout the recording session. In the case of the SNX experiments, the hippocampal window was also punctured. We then injected ∼50 nl of either SNX (10 µM) dissolved in sterile saline or sterile saline alone onto the distal apical dendritic region of CA1 (injection depth of ∼320 µm below the hippocampal surface), using the same procedure as described above for the virus injections. Subsequently, the hole was covered with Kwik-Cast, and the animal was allowed to recover for ∼45 minutes. Two-photon Ca^2+^ imaging proceeded then as usual. Notably, there was no difference in the licking or running behaviors between the standard experiments and those involving local pharmacology (Extended Data Figs. 2-3).

### Optogenetic perturbation of entorhinal cortex layer 3 activity

To preferentially manipulate EC3 activity, *loxP*-flanked Archaerhodopsin-T (ArchT)^21^ driven by a synapsin promoter was virally expressed by injecting AAVrg carrying the *loxP*-flanked ArchT-tdtomato payload into the area CA1 of pOxr1-cre mice (see above)^19^, which express Cre recombinase mostly in layer 3 neurons of the medial entorhinal cortex. The hippocampal window was implanted during the same surgery, and a fiber-containing ferrule was inserted into the EC. The ferrule contained a 200 µm core, 0.5 NA, multimode fiber (FP200ERT, Thorlabs) and was constructed using published techniques^43^. Approximately 21 days after virus injection, combined two-photon imaging and optogenetic experiments were performed. ArchT was activated using light pulses (maximal duration of 5 seconds, 594 nm, 40 Hz, sinusoidal pattern, Mambo laser, Cobalt, CA, USA), delivered through the optical fiber located in the EC. The mean laser power was 5-10 mW^22^ (measured each day before the recording in air, ∼0.5 cm from the tip of the fiber optic patch cable). As a control, the fluorescent protein tdtomato was virally expressed in pOxr1-Cre mice. These control mice were treated the same as the ArchT group.

To confirm an effect of the ArchT activation on EC3 activity, we performed in a group of mice (n=4) that expressed ArchT in EC3 local field potential (LFP) recordings in the entorhinal cortex. Glass electrodes (1.5-3.5 MΩ) were filled with 0.9% saline and mounted vertically on a micromanipulator (Luigs &Neumann). The LFP signal was monitored using an audio amplifier (Grass Technologies), while the electrode was advanced slowly through the brain with ∼0.5 psi of pressure. The LFP recording locations were ∼1.7 mm below the cortical surface. Once this depth was reached, we removed the pressure and started recordings. We alternated between control laps without and laps with ArchT activation. There was no randomization in the sequential ordering of control laps and laps with light application.

### Histology

Mice were transcardially perfused with phosphate-buffered saline (PBS) or Dulbecco’s PBS, followed by a 4% paraformaldehyde (PFA) solution. Extracted brains remained overnight in 4% PFA and were then rinsed twice and stored in PBS. Then, 50 μm-thick coronal or sagittal sections of paraformaldehyde-fixed brains were made and mounted on glass slides using Fluoromount mounting medium. All histological images were acquired on the ZEISS Axioscope, equipped with ZEN software.

### Data analysis

#### Ca^2+^ signal extraction and activity map generation

To extract somatic Ca^2+^ signals of CA1 pyramidal neurons, videos were motion-corrected using SIMA, regions of interest (ROIs) were manually drawn to include single neurons (using Image J), and calcium traces across time were extracted again using SIMA^44^. To extract axonal EC3 Ca^2+^ signals, the automatic motion correction and ROI detection algorithms of the Suite2P pipeline^45^ were used. The output was manually curated for both recording types, and ROIs with insufficient signal were removed. Only datasets, for which the motion correction was successful, were included in this study. Further analyses of CA1 and EC3 activity were then performed using custom functions written in MATLAB (Version 2019a). These included: 1) Conversion to Δf/f, where Δf/f was calculated as (F – F0)/F0, where F0 is the mode of the histogram of F; 2) In the case of the axonal Ca^2+^ data, a noise correlation analysis using a Pearson correlation coefficient threshold of 0.4-0.5 to identify ROIs that likely originate from the same axon/neuron (step-by-step procedure illustrated in Extended Data Fig. 5a-d). Ca^2+^ signals from ROIs belonging to a single axon were combined, and an average Ca^2+^ signal per axon was calculated, with ROIs weighted according to their size (=pixel number); 3) Detection of significant Ca^2+^ transients, i.e., transients larger than 3 * standard deviations of the noise (= baseline F values). We then produced Ca^2+^ activity maps across all spatial locations and laps for each CA1 pyramidal cell and EC3 axon, using only those recording epochs, during which the animal was running (velocity >2 cm/sec). These activity maps were generated by first dividing the length of the belt (= lap of 180 cm) into 50 spatial bins (3.6 cm each). For each spatial bin, the mean Δf/f was calculated. All Ca^2+^ activity maps were then smoothed using a 3 point boxcar, and for display purposes, aligned such that the opening of the valve (= reward delivery site) was either located in spatial bin 26 (Figs. 1-4, Extended Data Fig. 2, data recorded in environment A) or in spatial bin 24 (Fig. 5, Extended Data Figs. 6, 8, and 10, data recorded in environment B, or when environments A and B are compared). Visual stimulation and reward locations are marked by arrows or dashed lines in all figures. All recorded laps were included, except for the data presented in Figs. 4a-b and 5d-f as well as Extended Data Figs. 6a-f and 8d-e (analysis of stochastic firing properties of EC3 axons), where only laps 1-50 were used.

#### CA1 place cell identification

Many CA1 neurons were initially silent and acquired a place field suddenly during the recording sessions on days 0 or 1. Therefore, we first identified for each CA1 neuron a potential place field onset lap (‘induction lap’). A place field onset was defined as 1) a lap with a spatial bin with significant Ca^2+^ activity (greater than 3 * standard deviation of the noise) in lap X. 2) presence of spatial bins with significant Ca^2+^ activity in 2 out of the 5 following laps (lap X+1 to lap X+6). If more than 1 lap per neuron fit these criteria, we selected the first one, unless the field that was generated was weak and disappeared for more than 20 laps at some point during the recording. Only laps following the induction lap (= lap X) were used to determine whether a neuron was considered a place cell. Whether a CA1 neuron exhibited a spatially modulated field was defined 1) by the amount of spatial information its activity provided about the linear track position (>95% confidence interval of the shuffled spatial information values) and 2) by its reliability (significant activity in more than 30% of the laps following the induction lap). For each neuron, the spatial information SI was computed as described previously^46^:

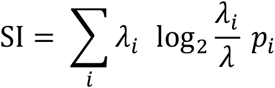

where p^i^ is the probability of occupancy in spatial bin i, λ_i_ is the mean activity level (Δf/f) while occupying bin i, and λ is the overall mean activity level (Δf/f). This value was compared to 100 shuffles of the activity (each shuffle was generated by circularly shifting the fluorescence trace by at least 500 frames, then dividing the fluorescence trace into six chunks and permuting their order). If the observed information value exceeded the 95% confidence interval of the shuffled information values, its field was considered spatially modulated. Neurons with no significant activity in any of the laps (‘silent neurons’) were not included in this analysis. The place field width was quantified as the number of consecutive spatial bins * 3.6 cm, for which the mean Δf/f exceeded 20% of the peak Δf/f value. Only one place field per neuron was included in our analyses.

#### Behavioral data quantification

Similar to the Ca^2+^ imaging data analysis, the running and licking behavioral maps were generated by first dividing the length of the belt (=lap of 180 cm) into 50 spatial bins (3.6 cm each). Then, the lick rate (licks per second, Hz) and the mean velocity (cm/sec) were calculated for each spatial bin.

#### Velocity correlation of EC3 axons

To categorize axons as significantly positively or negatively correlated with speed, the Pearson Correlation (matlab function ‘corr’) was calculated between the mean Δf/f Ca^2+^ activity and the mouse’s velocity (maps of 50 spatial bins per lap were used).

#### Computational Model

2000 two-state Markov Chains, 510 seconds in duration, were generated using the transition probabilities shown in the matrix in Extended Data Fig. 6f to simulate 50 laps each 10 seconds in duration with a time step of 0.1s (5100 time steps total). Each of these chains simulated the persistent firing activity of a single EC3 neuron, where state 0 was inactive and state 1 was active. The Markov chains were produced by randomly sampling numbers from a uniform distribution between 1 and 1000 on each time step. At each time step, a chain transitioned from inactive to active if the sampled number was less than or equal to (P_01_·100). Likewise, a chain transitioned from active to inactive if a randomly sampled number was less than or equal to (P_10_·100). This produced exponentially distributed active and inactive times with means (τ_on_ and τ_off_), as expected from the transition probabilities (Extended Data Fig. 7i). For example, the probability of a chain transitioning from inactive to active during one time step (Δt) is p=P_01_·Δt, giving a mean inactive time where ττ_inactive_ =Δt/p or 1/P_01_^47^. Each of the fifty 100 time-step sections of the chains were averaged and smoothed with a 3 point boxcar. The initial 100 points of each chain were not used to allow proper initialization. For the activity versus position plots, the average activity in each spatial bin was calculated using the actual mean running speed of the animals.

In addition to this set of chains using static or homogenous transition probabilities, we also used two other conditions. In these conditions, we were attempting to simulate the presence of two populations of chains, one that was purely homogenous without any changes in their transition probabilities and a second population that had transition probabilities that were sensitive to the environment. We used two pieces of the EC3 data to direct our manipulations. The first was the median cell-cell correlations, and the second was the spatial distribution of peak activity (Extended Data Figs. 6-8). Thus, for the uniform track used in environment A, we altered the transition probabilities in the same proportion of chains uniformly across the lap (Extended Data Fig. 7). For these conditions, at each of the 100 time steps, P_01_ was step increased from 0.04 to 0.24 for 1 s (10 time steps) in 14 chains for a total of 1400 chains in which the activation transition probability was adjusted. In the remaining 600 chains, P01 was not changed (homogenous condition). To simulate the nonuniform environment, we chose to manipulate the number of chains with increased P_01_ around 40 cm such that the final fraction of chains with peak activity around the light position was increased approximately three-fold (Extended Data Fig. 8). To do so, we used a similar procedure as above, except the number of chains with increased P_01_ around 40 cm was elevated according to the density plot in Extended Data Fig. 8a. The additional chains around the light stimulus were taken from the unmodulated pool, which was reduced to a total of 150 chains. While this alone increased the median cell-cell correlations, it was also necessary to slightly increase the activation transition probability in all chains (P_01_ stepped from 0.04 to 0.28 for 1 s) to approach the elevated median correlations observed in the data. To compare odd and even laps correlations, only a population of 1000 chains was used (but with all the same proportions) to simulate the experimental conditions more accurately.

These chains were used to predict the spatial plateau probability profile in a population of postsynaptic CA1 neurons. To do this, we randomly selected 100 of the 2000 chains and summed them (this represents 5% of the total “input” population). We chose 2000 because this is approximately the number of SLM synapses on CA1 pyramidal neurons, and 5% seemed like a reasonable fraction of active inputs. We have altered this number between 2.5 and 10% and found the results to be consistent between 5-10%. Next, feedforward inhibition was simulated simply as the sum of all 2000 chains scaled by the appropriate fraction (i.e., multiplied by 0.05), and this waveform was subtracted from the sum of the 100 “excitatory EC3 inputs”. This procedure was repeated 10,000 separate times to mimic postsynaptic integration in a large population of CA1 neurons. Finally, a threshold was set based on the observed fraction of the total CA1 population to generate new place fields during a session (20-25%). The fraction of “CA1 neurons” that crossed threshold (our proxy for plateau initiation probability) was calculated as the total number of threshold crossings in 30 spatial bins divided by the total number of “neurons” (10,000) using the actual running speed profile of the animals or a constant speed profile of 18 cm/s to determine the dwell time in each bin.

#### Statistical methods

The exact sample size (n) for each experimental group is indicated in the figure legend or in the main text. No statistical methods were used to predetermine sample sizes, but our sample sizes are similar to those reported in previous publications^6,24^. In some cases, when data distribution was assumed, but not formally tested, to be normal, data were analyzed using two-tailed paired or unpaired *t*-tests, as stated in the text or figure legends. Data were analyzed automatically without consideration of trial conditions or experimental groups. Experiments and data analyses were not randomized and not performed blind to the experimental conditions. If not otherwise indicated in the figure, data are shown as mean ± SEM.

#### Data Availability

The data that support the findings of this study are available from the corresponding author upon request.

#### Code Availability

The code that supports the findings of this study are available from the corresponding author upon request.

## References

1 Abbott, L. F. & Nelson, S. B. Synaptic plasticity: taming the beast. Nat Neurosci 3 Suppl, 1178–1183, (2000).

2 Martin, S. J., Grimwood, P. D. & Morris, R. G. Synaptic plasticity and memory: an evaluation of the hypothesis. Annu Rev Neurosci 23, 649–711, (2000).

3 Dupret, D., O’Neill, J., Pleydell-Bouverie, B. & Csicsvari, J. The reorganization and reactivation of hippocampal maps predict spatial memory performance. Nat Neurosci 13, 995–1002, (2010).

4 Hollup, S. A., Molden, S., Donnett, J. G., Moser, M. B. & Moser, E. I. Accumulation of hippocampal place fields at the goal location in an annular watermaze task. J Neurosci 21, 1635–1644, (2001).

5 Turi, G. F., Li, W. K., Chavlis, S. et al. Vasoactive Intestinal Polypeptide-Expressing Interneurons in the Hippocampus Support Goal-Oriented Spatial Learning. Neuron 101, 1150–1165 e1158, (2019).

6 Zaremba, J. D., Diamantopoulou, A., Danielson, N. B. et al. Impaired hippocampal place cell dynamics in a mouse model of the 22q11.2 deletion. Nat Neurosci 20, 1612–1623, (2017).

7 Caporale, N. & Dan, Y. Spike timing-dependent plasticity: a Hebbian learning rule. Annu Rev Neurosci 31, 25–46, (2008).

8 Mehta, M. R., Quirk, M. C. & Wilson, M. A. Experience-dependent asymmetric shape of hippocampal receptive fields. Neuron 25, 707–715, (2000).

9 Moore, J. J., Cushman, J. D., Acharya, L., Popeney, B. & Mehta, M. R. Linking hippocampal multiplexed tuning, Hebbian plasticity and navigation. Nature, (2021).

10 Dana, H., Chen, T. W., Hu, A. et al. Thy1-GCaMP6 transgenic mice for neuronal population imaging in vivo. PLoS One 9, e108697, (2014).

11 Bittner, K. C., Milstein, A. D., Grienberger, C., Romani, S. & Magee, J. C. Behavioral time scale synaptic plasticity underlies CA1 place fields. Science 357, 1033–1036, (2017).

12 Bittner, K. C., Grienberger, C., Vaidya, S. P. et al. Conjunctive input processing drives feature selectivity in hippocampal CA1 neurons. Nat Neurosci 18, 1133–1142, (2015).

13 Grienberger, C., Chen, X. & Konnerth, A. NMDA receptor-dependent multidendrite Ca(2+) spikes required for hippocampal burst firing in vivo. Neuron 81, 1274–1281, (2014).

14 Grienberger, C., Milstein, A. D., Bittner, K. C., Romani, S. & Magee, J. C. Inhibitory suppression of heterogeneously tuned excitation enhances spatial coding in CA1 place cells. Nat Neurosci 20, 417–426, (2017).

15 Takahashi, H. & Magee, J. C. Pathway interactions and synaptic plasticity in the dendritic tuft regions of CA1 pyramidal neurons. Neuron 62, 102–111, (2009).

16 Zhao, X., Wang, Y., Spruston, N. & Magee, J. C. Membrane potential dynamics underlying context-dependent sensory responses in the hippocampus. Nat Neurosci 23, 881–891, (2020).

17 Megias, M., Emri, Z., Freund, T. F. & Gulyas, A. I. Total number and distribution of inhibitory and excitatory synapses on hippocampal CA1 pyramidal cells. Neuroscience 102, 527–540, (2001).

18 Steward, O. & Scoville, S. A. Cells of origin of entorhinal cortical afferents to the hippocampus and fascia dentata of the rat. J Comp Neurol 169, 347–370, (1976).

19 Suh, J., Rivest, A. J., Nakashiba, T., Tominaga, T. & Tonegawa, S. Entorhinal cortex layer III input to the hippocampus is crucial for temporal association memory. Science 334, 1415–1420, (2011).

20 Tervo, D. G., Hwang, B. Y., Viswanathan, S. et al. A Designer AAV Variant Permits Efficient Retrograde Access to Projection Neurons. Neuron 92, 372–382, (2016).

21 Chow, B. Y., Han, X., Dobry, A. S. et al. High-performance genetically targetable optical neural silencing by light-driven proton pumps. Nature 463, 98–102, (2010).

22 Li, N., Chen, S., Guo, Z. V. et al. Spatiotemporal constraints on optogenetic inactivation in cortical circuits. Elife 8, (2019).

23 Campbell, M. G., Ocko, S. A., Mallory, C. S. et al. Principles governing the integration of landmark and self-motion cues in entorhinal cortical codes for navigation. Nat Neurosci 21, 1096–1106, (2018).

24 Cholvin, T., Hainmueller, T. & Bartos, M. The hippocampus converts dynamic entorhinal inputs into stable spatial maps. Neuron 109, 3135–3148 e3137, (2021).

25 Hardcastle, K., Maheswaranathan, N., Ganguli, S. & Giocomo, L. M. A Multiplexed, Heterogeneous, and Adaptive Code for Navigation in Medial Entorhinal Cortex. Neuron 94, 375–387 e377, (2017).

26 Tang, Q., Ebbesen, C. L., Sanguinetti-Scheck, J. I. et al. Anatomical Organization and Spatiotemporal Firing Patterns of Layer 3 Neurons in the Rat Medial Entorhinal Cortex. J Neurosci 35, 12346–12354, (2015).

27 Egorov, A. V., Hamam, B. N., Fransen, E., Hasselmo, M. E. & Alonso, A. A. Graded persistent activity in entorhinal cortex neurons. Nature 420, 173–178, (2002).

28 Hahn, T. T., McFarland, J. M., Berberich, S., Sakmann, B. & Mehta, M. R. Spontaneous persistent activity in entorhinal cortex modulates cortico-hippocampal interaction in vivo. Nat Neurosci 15, 1531–1538, (2012).

29 Tahvildari, B., Fransen, E., Alonso, A. A. & Hasselmo, M. E. Switching between “On” and “Off” states of persistent activity in lateral entorhinal layer III neurons. Hippocampus 17, 257–263, (2007).

30 Enikolopov, A. G., Abbott, L. F. & Sawtell, N. B. Internally Generated Predictions Enhance Neural and Behavioral Detection of Sensory Stimuli in an Electric Fish. Neuron 99, 135–146 e133, (2018).

31 Milstein, A. D., Li, Y., Bittner, K. C. et al. Bidirectional synaptic plasticity rapidly modifies hippocampal representations independent of correlated activity. bioRxiv, (2020).

32 Royer, S., Zemelman, B. V., Losonczy, A. et al. Control of timing, rate and bursts of hippocampal place cells by dendritic and somatic inhibition. Nat Neurosci 15, 769–775, (2012).

33 Kaufman, A. M., Geiller, T. & Losonczy, A. A Role for the Locus Coeruleus in Hippocampal CA1 Place Cell Reorganization during Spatial Reward Learning. Neuron 105, 1018–1026 e1014, (2020).

34 Bloss, E. B., Cembrowski, M. S., Karsh, B. et al. Single excitatory axons form clustered synapses onto CA1 pyramidal cell dendrites. Nat Neurosci 21, 353–363, (2018).

35 Williams, S. R. & Fletcher, L. N. A Dendritic Substrate for the Cholinergic Control of Neocortical Output Neurons. Neuron 101, 486–499 e484, (2019).

36 Labarrera, C., Deitcher, Y., Dudai, A. et al. Adrenergic Modulation Regulates the Dendritic Excitability of Layer 5 Pyramidal Neurons In Vivo. Cell Rep 23, 1034–1044, (2018).

37 Buesing, L., Bill, J., Nessler, B. & Maass, W. Neural dynamics as sampling: a model for stochastic computation in recurrent networks of spiking neurons. PLoS Comput Biol 7, e1002211, (2011).

38 Gershman, S. J., Vul, E. & Tenenbaum, J. B. Perceptual multistability as Markov chain Monte Carlo inference. Advances in Neural Information Processing Systems (NIPS), (2009).

39 Bengio, Y. Deriving differential target propagation from iterating approximate inverses. arXiv preprint 2007.15139, (2020).

40 Lillicrap, T. P., Santoro, A., Marris, L., Akerman, C. J. & Hinton, G. Backpropagation and the brain. Nat Rev Neurosci 21, 335–346, (2020).

41 Kostadinov, D., Beau, M., Blanco-Pozo, M. & Hausser, M. Predictive and reactive reward signals conveyed by climbing fiber inputs to cerebellar Purkinje cells. Nat Neurosci 22, 950–962, (2019).

42 Raymond, J. L. & Medina, J. F. Computational Principles of Supervised Learning in the Cerebellum. Annu Rev Neurosci 41, 233–253, (2018).

## References

43 Sparta, D. R., Stamatakis, A. M., Phillips, J. L. et al. Construction of implantable optical fibers for long-term optogenetic manipulation of neural circuits. Nat Protoc 7, 12–23, (2011).

44 Kaifosh, P., Zaremba, J. D., Danielson, N. B. & Losonczy, A. SIMA: Python software for analysis of dynamic fluorescence imaging data. Front Neuroinform 8, 80, (2014).

45 Pachitariu, M., Stringer, C., Dipoppa, M. et al. Suite2p: beyond 10,000 neurons with standard two-photon microscopy. BioRxiv, (2017).

46 Gauthier, J. L. & Tank, D. W. A Dedicated Population for Reward Coding in the Hippocampus. Neuron 99, 179–193 e177, (2018).

47 Colquhoun, D. & Hawkes, A. G. The interpretation of single channel recordings. Microelectrode Techniques, (1987).

