## Supplemental Figures for "Entorhinal cortex directs learning-related changes in CA1 representations"

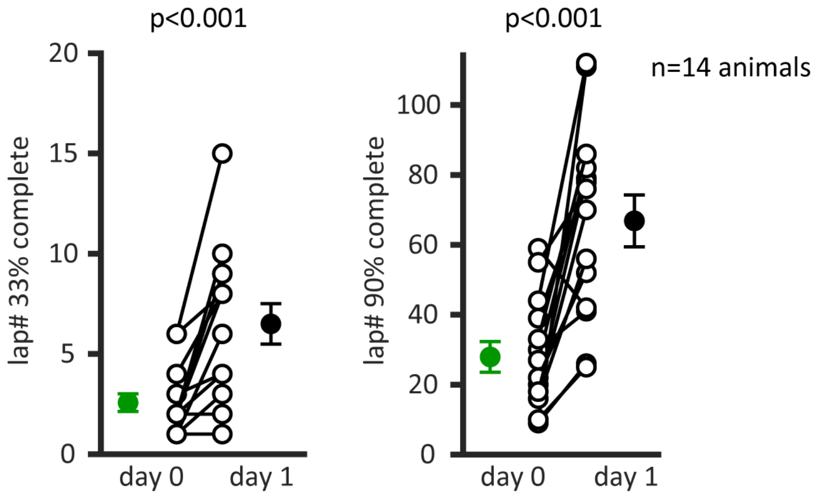


**Extended Data Figure 1: Time course of the CA1 place cell representation development.** Shown are the numbers of laps required to build 33% (left) and 90% (right) of the CA1 place cell representation on day 0 and day 1 (33%: n=14; two-tailed paired *t*-test, day 0 vs. day 1, $p<0.001$; 90%: n=14; two-tailed paired *t*-test, day 0 vs. day 1, $p<0.001$. The open circles show individual animals, the filled circles the mean. Data are shown as mean +/- SEM.

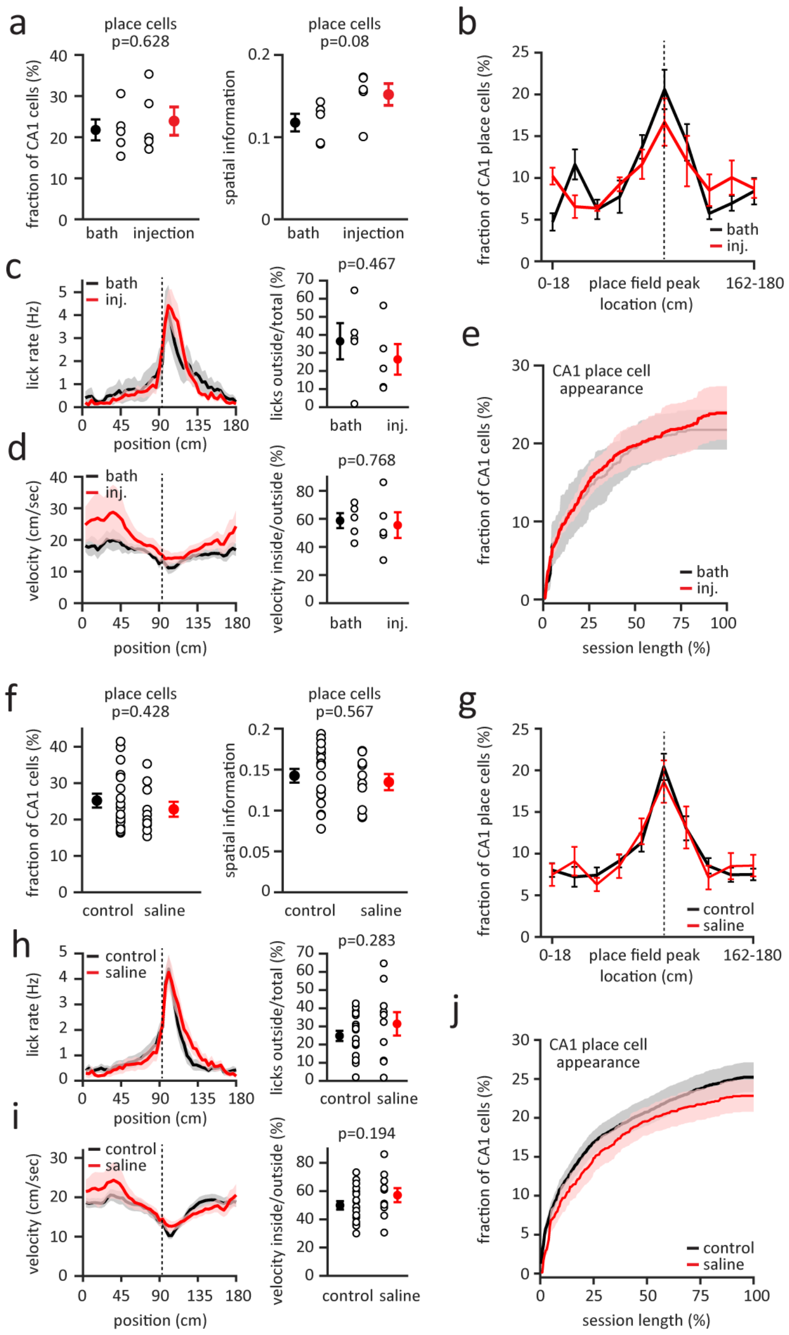


**Extended Data Figure 2: Bath application and injection of sterile saline 0.9% do not impact hippocampal activity and behavior. a-e**, Comparison of bath application (black, n=5 animals) and injection (red, n=5 animals) of sterile saline 0.9% in environment A. **a,** Left: Fraction of CA1 cells that are spatially modulated (two-tailed unpaired *t*-test, bath vs. injection, p=0.628). Right: Mean place cell spatial information content per animal (two-tailed unpaired *t*-test, bath vs. injection, p=0.08). **b,** Fraction of CA1 place cells as a function of place field peak location. The track is divided into ten spatial bins of 18 cm. **c,** Left. Mean lick rates for bath application and injection of sterile saline 0.9%. Right: Mean number of licks outside the reward zone (from 14 cm before to 36 cm after the reward) divided by the total number of licks (two-tailed unpaired *t*-test, bath vs. injection, p=0.467). **d,** Left: Mean running profile for bath application and injection of sterile saline 0.9%. Right: Mean velocity inside the reward zone divided by mean velocity outside the reward zone (two-tailed unpaired *t*-test, bath vs. injection, p=0.768). **e,** Time course of CA1 place cell appearance for bath application and injection of sterile saline 0.9%. In panels a, c, d the open circles show individual animals, the filled circles the mean. **f-j,** Same as **a-e** for comparing control condition (black, no saline, n=18 animals) and sterile saline 0.9% application via bath application and injection (red, n=10 animals) in environment A. Black dashed lines depict the reward location. Data are shown as mean +/- SEM.

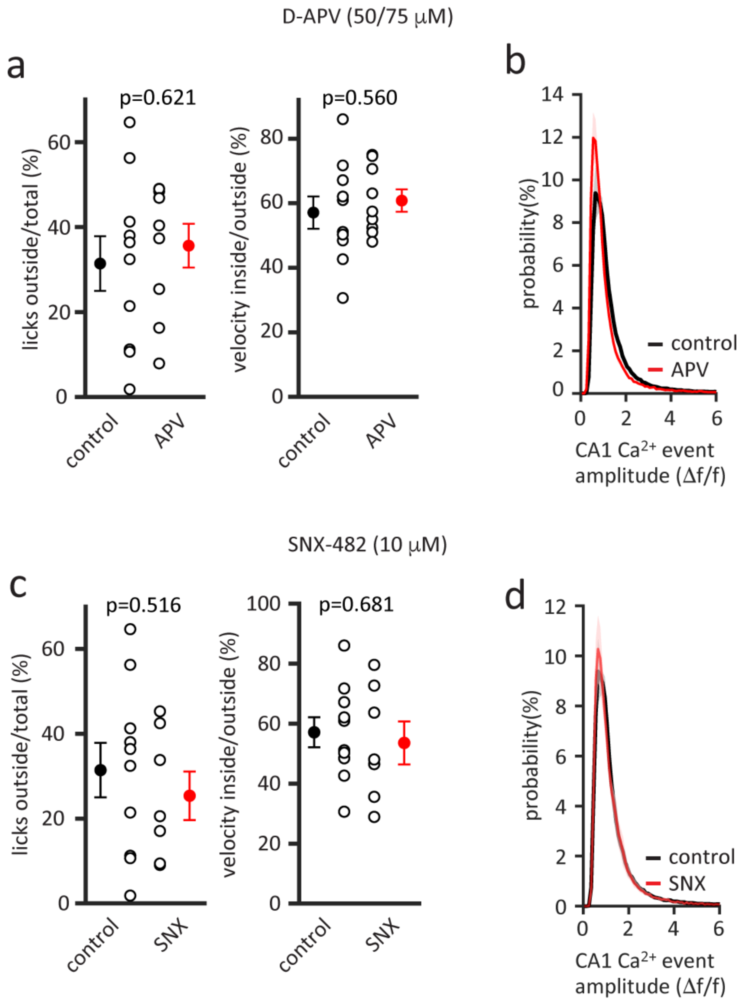


**Extended Data Figure 3: Effect of locally applied BTSP blockers on behavior and CA1 $Ca^{2+}$ event amplitude. a-b,** Effect of NMDA receptor antagonist, D-APV (50/75 μM). Black: Control (n=10 animals). Red: APV (n=8 animals). **a,** Left. Mean number of licks outside the reward zone (from 14 cm before to 36 cm after the reward) divided by the total lick number. Right: Mean velocity inside the reward zone divided by mean velocity outside the reward zone. The open circles show individual animals, the filled circles the mean. **b,** Distribution of CA1 $Ca^{2+}$ event amplitudes recorded. **c-d,** Effect of $Ca^{2+}$ channel antagonist, SNX-482 (10 μM). Black: Control (n=10 animals). Red: SNX (n=7 animals). Panels same as **a-b**. Two-tailed unpaired *t*-test were used, and data are shown as mean +/- SEM.

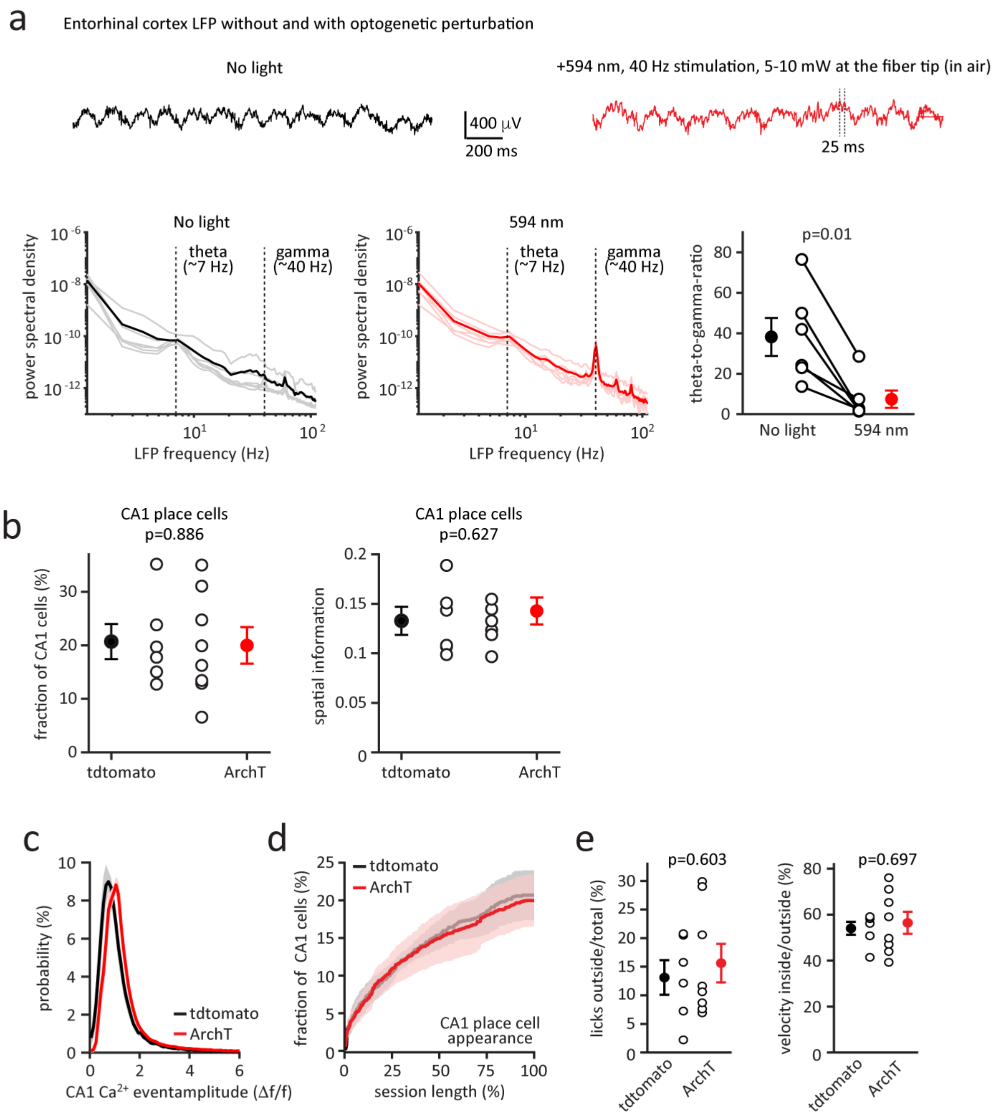


**Extended Data Figure 4: Optogenetic perturbation of entorhinal cortex layer 3 neurons via Archaerhodopsin T (ArchT).** **a,** Entorhinal cortex LFP without (black) or with (red) optogenetic perturbation via ArchT expression in layer 3 (n=6 sessions from 4 animals). Top. Raw LFP signal recorded in the same animal. Bottom: Power spectral density analysis (thin lines: individual sessions; thick lines: mean) and theta-to-gamma-ratio. **b-e,** Basic CA1 place cell features and the behavior are indistinguishable in tdtomato control (black, n=6) and ArchT (red, n=8) animals. **b,** Left: Fraction of CA1 cells that are spatially modulated (two-tailed unpaired *t*-test, p=0.886). Right: Mean place cell spatial information content per animal (two-tailed unpaired *t*-test, p=0.627). **c,** Distribution of CA1 $Ca^{2+}$ event amplitudes recorded. **d,** Time course of CA1 place cell appearance for tdtomato control and ArchT animals. **e,** Left: Mean number of licks outside the reward zone (from 14 cm before to 36 cm after the reward) divided by the total number of licks (two-tailed unpaired *t*-test, p=0.603). Right: Mean velocity inside the reward zone divided by mean velocity outside the reward zone (two-tailed unpaired *t*-test, p=0.697). In panels a, b and e, the open circles show individual animals, the filled circles the mean. Data are shown as mean +/- SEM.

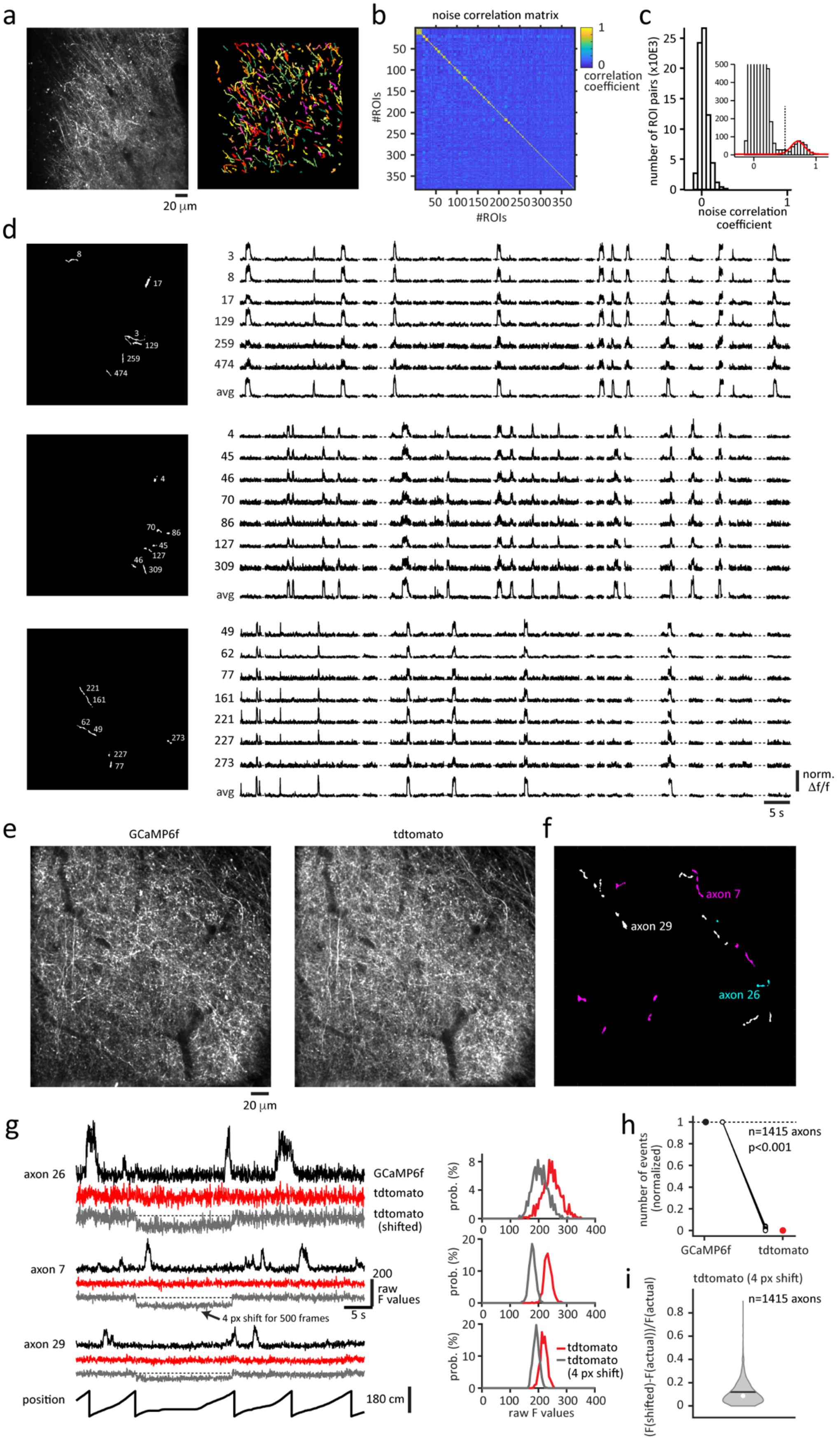


**(caption on the next page)**

**Extended Data Figure 5: Entorhinal cortex layer 3 axon imaging. a-c,** Analysis pipeline to identify ROIs that belong to the same axon. Left. Single-plane, two-photon, time-averaged image showing expression of GCaMP6f in EC3 axons. Right. All axonal regions of interest (ROIs), as identified by Suite2p. Colors are assigned randomly. **b,** Noise correlation matrix for all axonal ROIs identified in this animal (n=369). ROIs are sorted so that highly correlated ROIs are clustered. **c,** Histogram showing the noise correlation coefficient distribution for all ROI pairs from this animal. Inset shows magnified view of second, distinct, histogram peak (red line depicts gaussian fit of this second peak). All ROIs with a noise correlation coefficient value > 0.5 were assumed to belong to the same axon and subsequently combined into a single compound ROI. **d,** 3 example axons. Left: white areas depicting individual ROIs, which are assumed to belong to a single axon. Right. Normalized Δf/f traces for these ROIs (numbers correspond to the masks on the left) and for the resulting single axon (avg). The gaps represent epochs during which $Ca^{2+}$ signal was not recorded (e.g., because the animal was stationary). **e-i**, Simultaneous imaging of GCaMP6f and tdtomato in EC3 axons as control for z-motion. **e,** Single-plane, two-photon, time-averaged image showing expression of the GCaMP6f (green channel) and tdtomato (red channel) in EC3 axons. **f,** Colored areas depict three individual axons, as identified by the noise correlation analysis. **g,** Left. Raw fluorescence traces (black: GCaMP6f, red: tdtomato) for the three axons. The grey traces are obtained by shifting all ROIs belonging to the axon for 500 frames by 4 px in the x-dimension. The position signal at the bottom indicates epochs of varying running speeds. Regardless of the animal's velocity, the tdtomato signal remains stable. Right: Raw F value histograms of the 500-frame period where the axonal ROIs are shifted. **h,** Number of events (normalized) per axon (n=1415 axons from 14 animals). The open circles show individual animals, the filled circles the mean. **i,** Difference between recorded raw tdtomato fluorescence values and the fluorescence values when ROIs are artificially shifted by 4 pixels in the x dimension. The difference is shown as a fraction of the actual value. The white dot marks the median, the black line the mean of the distribution. A 4-pixel shift (~2 μm) would have caused a detectable (~10%) change in the tdtomato fluorescence. If not otherwise indicated, data are shown as mean +/- SEM.

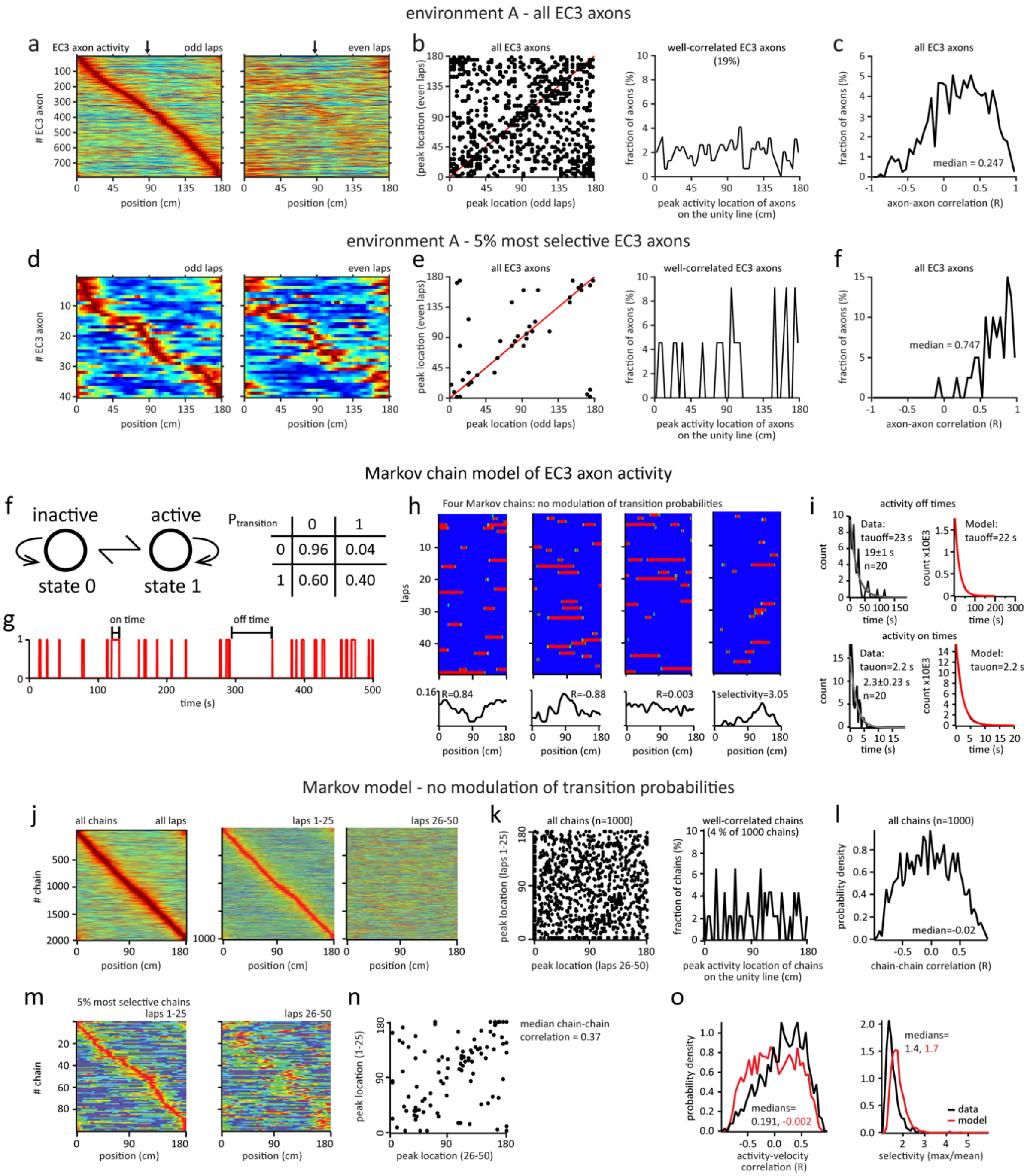


**Extended Data Figure 6: Modeling of EC3 axon activity.** **a-f,** Characterization of EC3 axonal activity, recorded in environment A. **a,** Normalized mean $\Delta f/f$ across space for all entorhinal cortex layer 3 (EC3) axons recorded in environment A (n=792 axons in 7 animals). Color plots for odd and even laps are shown separately. EC3 axons are ordered according to their peak location during the odd laps. **b,** Left: Scatter plot showing axonal activity peak locations for averages made from odd and even laps. The unity line is depicted in red. Right: Histogram of peak activity locations of well-correlated axons. Peak locations on even trials were within 10 cm of odd trials. **c,** Distribution of Pearson's correlation coefficients for axon-axon comparisons from data in **b**. **d-f,** Same as panels **a-c**, but only the 5% most selective axons are included. **f-o,** Markov chain model of EC3 axon activity. **f,** Individual EC3 axon activity was modeled as a two-state Markov chain with base transition probabilities as shown in the matrix. Each chain, 2000 in total, was generated as transitions from 0 (inactive) to 1 (active) for 50 laps with a single lap duration of 10 s. **g,** Representative chain showing the transitions used to calculate active (on) times and inactive (off) times. **h,** Activity heat maps for 50 laps (10 s each) for four different chains. From left to right, chain showing high positive velocity correlation, chain showing high negative velocity correlation, chain showing no correlation and chain showing strong selectivity (maximum amplitude/mean amplitude of average trace. Average traces shown below. **i,** Activity times are exponentially distributed for real EC3 single axons (left plots, black, 20 axons with highest number of events) and the population of model chains (right plots, red, all chains). **j,** Chains with static (unmodulated) transition probabilities. From left to right: heat maps of peak scaled mean chain activity for 50 laps from population of 2000 chains plotted in space. Average activity for the first 25 laps for subset of 1000 chains. Average activity for the last 25 laps for same chains. **k,** Left: plot of peak locations for average activity from first 25 laps versus last 25 laps. Right: Density of well-correlated chains (chains whose peak locations were within 10 cm in laps 1-25 and 26-50). **l,** Distribution of Pearson's correlation coefficients for chain-chain comparisons from data in **k**. **m-n,** Same as panels **j-k**, but only the 5% most selective chains are included. **o,** Distributions of activity-velocity correlation coefficients (left) and selectivity indices (right) for population of EC3 axons (black) and unmodulated model data (red). Medians are listed.

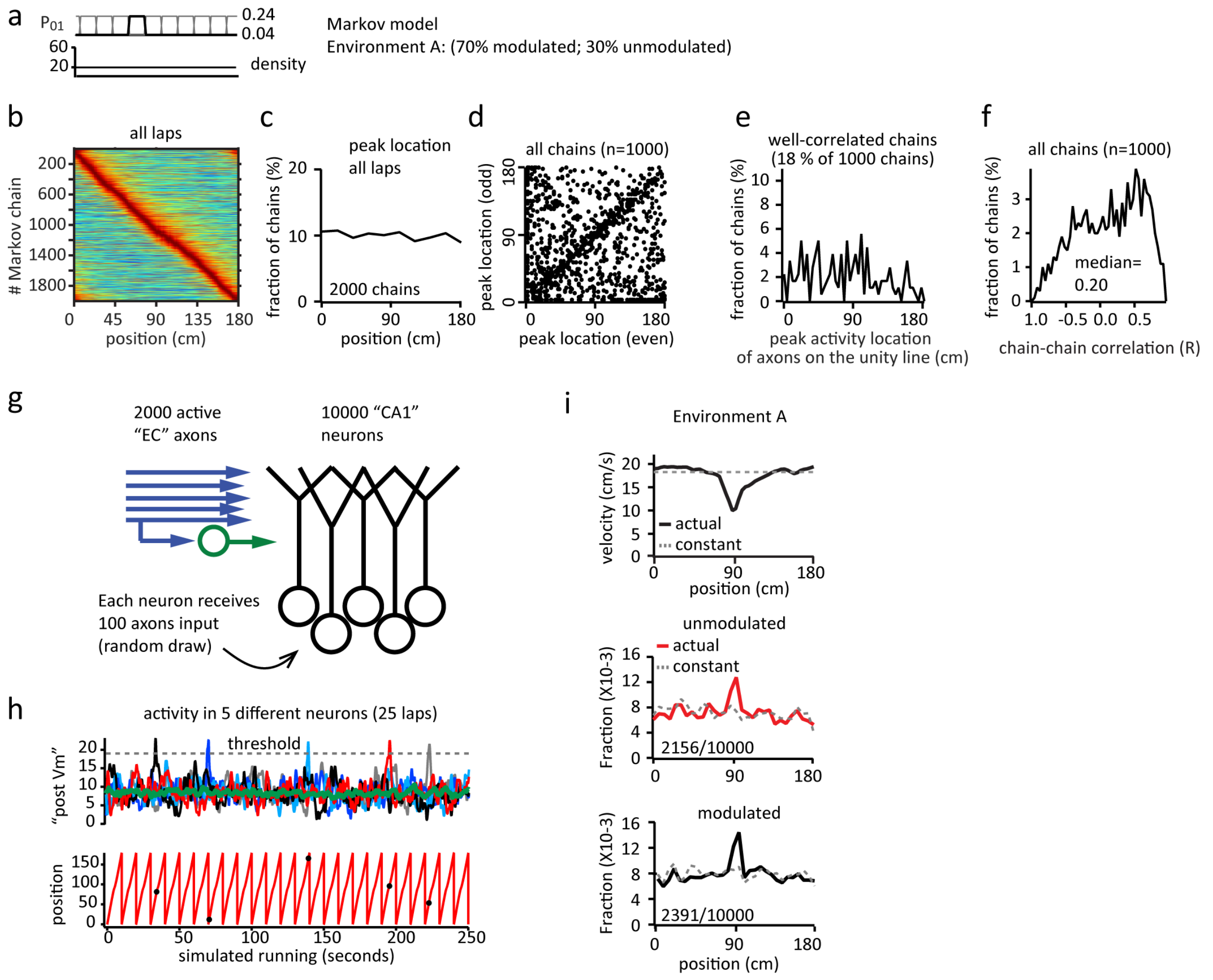


**Extended Data Figure 7: Analysis of the Markov chain model of EC3 axon activity and predictions for environment A. a,** Transition probabilities and chain density, where $P_{0,1}$ was modulated in 1400 chains by a 1 second step that moved iteratively across the lap. **b,** Heat maps of normalized mean chain activity for 50 laps from population of 2000 chains plotted in space. **c,** Fraction of Markov chains as a function of their peak activity location. **d,** Plot of peak locations for average activity from odd versus even laps. **e,** Histogram of peak activity locations of well-correlated chain. Peak locations on even trials were within 10 cm of odd trials. **f,** Distribution of Pearson's correlation coefficients for chain-chain comparisons from data in **d**. **g,** Model of impact on postsynaptic neurons. One hundred (5%) 50-lap chains are randomly sampled from the total of two thousand and summed 10,000 separate times to simulate postsynaptic summation in a population of 10,000 CA1 dendrites. In addition, an average activity from all 2000 (scaled appropriately: 0.05x) was subtracted from each summed chain to mimic feedforward inhibition. **h,** The activity in five representative postsynaptic neurons (100 summed chains) is shown for 25 consecutive laps. Inhibitory trace (green). Threshold, set so that 20-25% of postsynaptic neurons cross, is demarcated by dashed line. Below is the position of the mouse from actual data (black circles indicate location in space of threshold crossing). **i,** Plots simulate environment A showing from top to bottom: actual running speed profile in space (black) and a trace simulating a constant running speed (gray dashed). The fraction of postsynaptic neurons with threshold crossings for unmodulated condition and modulated condition vs. position on the track.

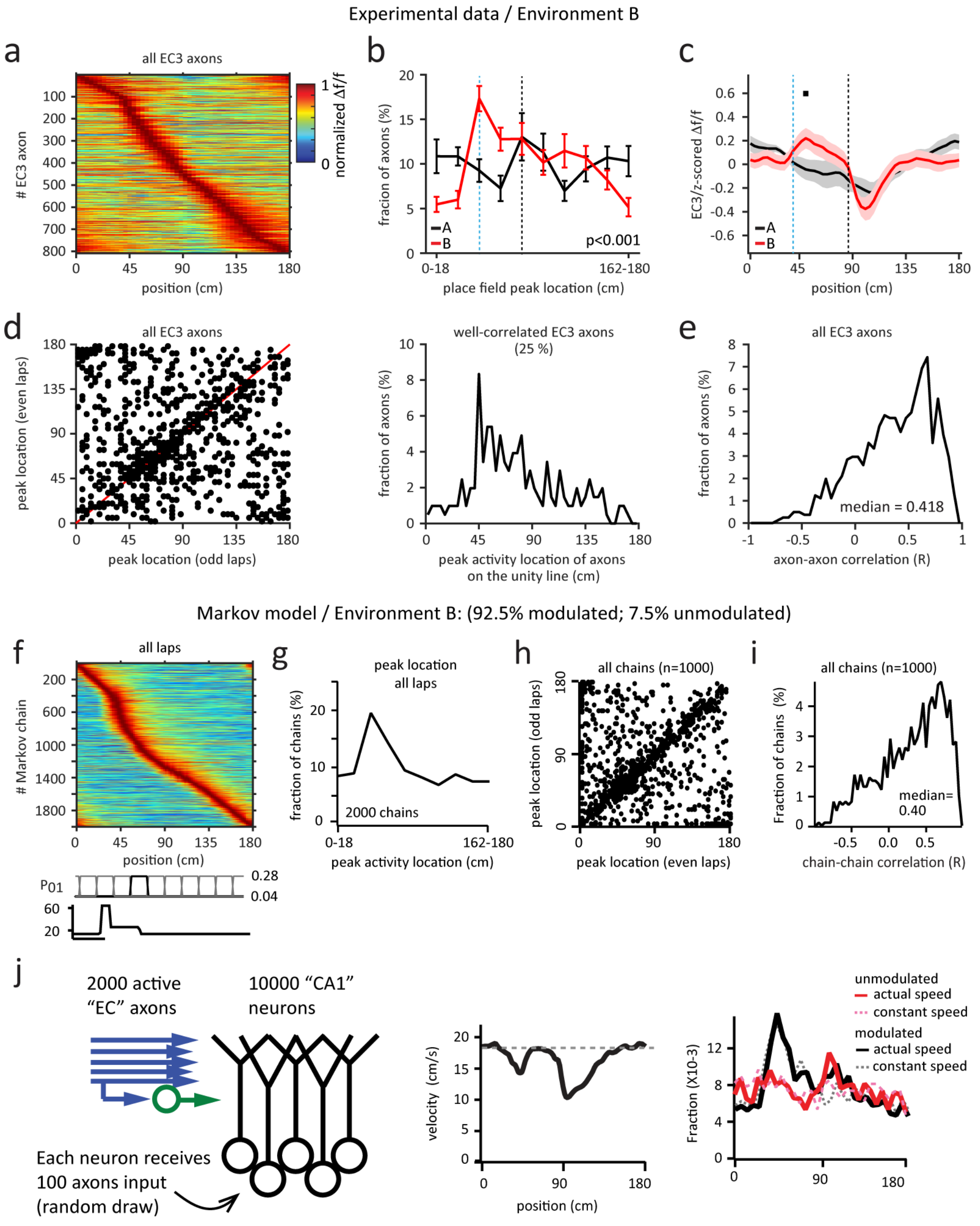


**Extended Data Figure 8: Analysis of EC3 axon activity in environment B and the predictions of the Markov chain model.** **a-e** Characterization of EC3 axonal firing in environment B. **a,** Normalized mean Δf/f across space for EC3 axons (n=808, n=8 animals) in env B. **b,** Fraction of EC3 axons as a function of their peak activity location (A: n=7, black; B: n=8, red; chi-square test, $p<0.001$). **c,** z-scored mean Δf/f of all EC3 axons. Black bars indicate locations with $p<0.05$ (unpaired two-tailed *t*-test). **d,** Left: Plot of peak locations for average activity from even laps versus odd laps. Right: Histogram of peak activity locations of well-correlated axons. Peak locations on even trials were within 10 cm of odd trials. **e,** Distribution of Pearson's correlation coefficients for axon-axon comparisons from data in **d**. **f-j,** Modeling of EC3 axon activity in environment B. **f,** Heat maps of normalized mean chain activity for 50 laps from population of 2000 chains plotted in space. **g,** Fraction of Markov chains as a function of their peak activity location. **h,** plot of peak locations for average activity from odd versus even laps. **i,** Distribution of Pearson's correlation coefficients for chain-chain comparisons from data in **h**. **j,** Left: Model of impact on postsynaptic neurons. Middle: actual running speed profile in space in environment B. Right: The fraction of postsynaptic neurons with threshold crossings for unmodulated condition and modulated condition vs position on the track. Blue dashed lines depict the light onset, black dashed lines the reward location. If not indicated otherwise, data are shown as mean +/- SEM.

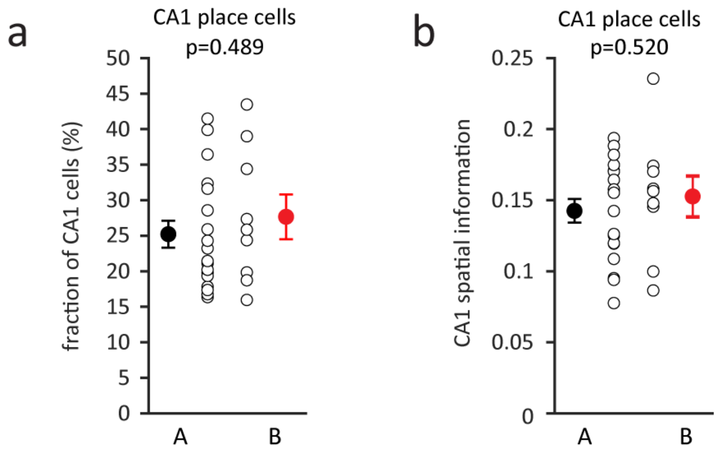


**Extended Data Figure 9: Basic CA1 place cell features are similar in environments A (somatosensory cues on the belt, n=18 animals) and B (no cues on the belt, visual stimulus only, n=9 animals). a,** Fraction of CA1 cells that are spatially modulated (two-tailed unpaired *t*-test, p=0.489). **b,** Mean place cell spatial information content per animal (two-tailed unpaired *t*-test, p=0.520). The open circles show individual animals, the filled circles the mean. Data are shown as mean +/- SEM.

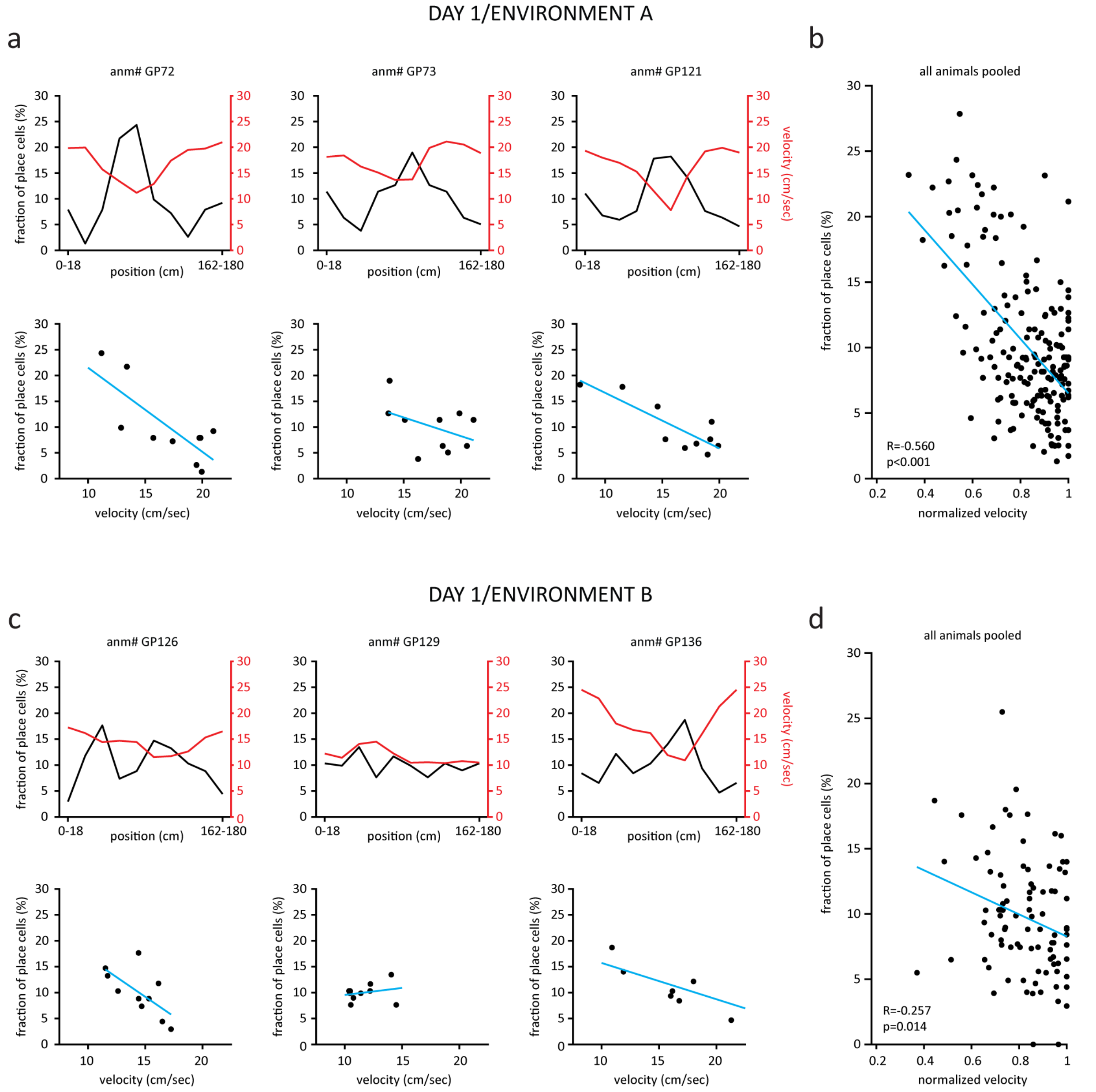


**Extended Data Figure 10: CA1 place cell density and the animals' velocity are inversely correlated. a,** Three example animals in environment A. Top: Fraction of CA1 place cells (black)/place cell density and the animal's mean velocity (red) as a function of location. Black and red y-axes apply, respectively. The track is divided into ten spatial bins of 18 cm. Bottom: Scatter plots showing the relationship between the CA1 place cell density and the animal's velocity at the same location. Each dot represents one spatial bin of 18 cm. Data points fit by linear equation (blue line). **b,** Scatter plot showing the relationship between the CA1 place cell density and the normalized velocity at the same location. Each dot represents one spatial bin of 18 cm. Data points pooled across all animals (n=18) fit by linear equation (blue line). **c-d,** Same as **a-b** for environment B (n=9 animals).

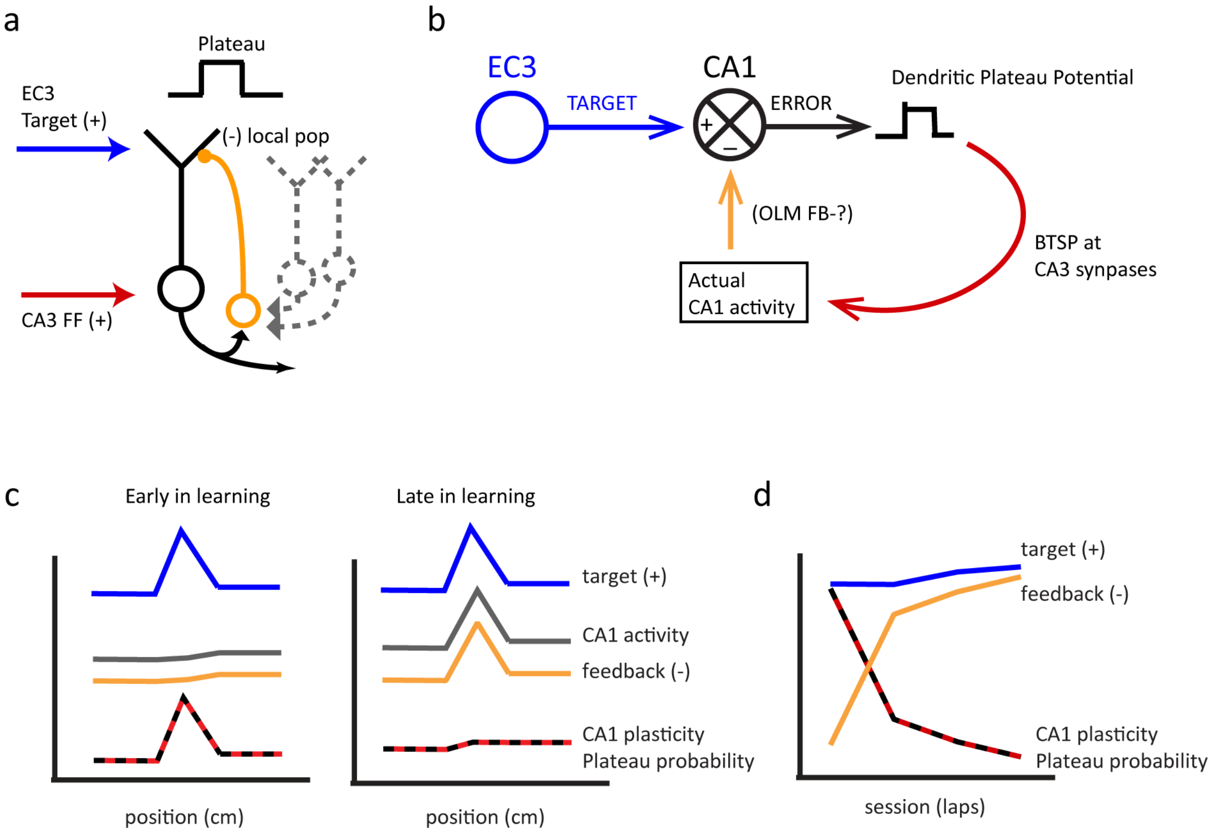


**Extended Data Figure 11: Hypothesized network scheme for learning in CA1. a,** CA1 circuit elements involved include CA1 pyramidal neurons (black solid and gray dashed), excitatory input from EC3 innervating the distal apical tuft regions and functioning as a target input (blue), excitatory input from CA3 whose synaptic weights are adjusted during learning innervates the perisomatic regions (red), OLM feedback inhibitory interneurons bring a copy of regional CA1 activity to the apical tuft (yellow). **b,** EC3 provides each individual CA1 neuron with a desired or target activity pattern that is compared in the distal apical dendrites with a representation of the actual pattern of population activity that could be provided by OLM-Interneurons. An excess excitation (mismatch) will increase the probability of dendritic $Ca^{2+}$ plateau potential initiation. The plateau functions as a local error signal in each CA1 cell that drives BTSP at CA3 feedforward excitatory inputs (the learning pathway) and shape the firing of each CA1 cell accordingly. This will change CA1 population activity that eventually feeds back to the comparator in the apical tuft. **c,** Spatial profiles of hypothesized signals. During the initial phase of learning there is an increased EC3 input (blue) around the reward site at the middle of the track that leads to increased plateau initiation and increased synaptic plasticity (red/black) there. Later in the session, the CA1 population (grey) shows an elevated activity near the middle of the track, which drives increased feedback inhibition (orange) that reduces plateau initiation and CA1 plasticity (red/black) even though the target input from the EC (blue) is unchanged. **d,** The changes in the signal amplitudes plotted across the session.
